# Cellular-resolution mapping uncovers spatial adaptive filtering at the cerebellum input stage

**DOI:** 10.1101/2020.03.14.991794

**Authors:** Casali Stefano, Tognolina Marialuisa, D’Angelo Egidio

## Abstract

Long-term synaptic plasticity, in the form of either potentiation or depression (LTP or LTD), is thought to provide the substrate for adaptive computations in brain circuits. Although molecular and cellular processes of plasticity have been clarified to a considerable extent at individual synapses, very little is known about the spatiotemporal organization of LTP and LTD in local microcircuits. Here, we have combined multi-spot two-photon laser microscopy and realistic modeling to map the distribution of plasticity in multi-neuronal units of the cerebellar granular layer activated by stimulating an afferent mossy fiber bundle. The units, composed by ~300 active neurons connected to ~50 glomeruli, showed potentiation concentrated in the core and depression in the periphery. This plasticity was effectively accounted for by an NMDA receptor and calcium-dependent induction rule and was regulated by local microcircuit mechanisms in the inhibitory Golgi cell loops. The organization of LTP and LTD created effective spatial filters tuning the time-delay and gain of spike retransmission at the cerebellum input stage and provided a plausible basis for the spatiotemporal recoding of input spike patterns anticipated by the motor learning theory.

## Introduction

Long-term synaptic plasticity (LTP and LTD) is usually investigated as a single synapse phenomenon under the assumption that multiple independent changes will eventually shape local field responses and brain computations^1–4^. In fact, individual neurons are integrated into local microcircuits, which form effective computational units like the cortical microcolumns ^5,6^ or the cerebellar microzones^7^. It is therefore conceivable that, like neuronal activity, also LTP and LTD are spatially coordinated within local neuronal assemblies that we will call “units”. However, the fine-grain organization of changes in such units remains largely unknown despite the potential impact it might have on microcircuit computation.

The cerebellum has long been thought to perform two main serial operations, i.e. spatiotemporal recoding of input signals in the granular layer followed by pattern recognition in the molecular layer^8,9^. The nature of these operations has been investigated in some details, showing that single granule cells can actually regulate the gain and timing of spike emission under control of the local inhibitory network formed by Golgi cell interneurons and of long-term synaptic plasticity at the mossy fiber – granule cell synapse^10–12^. But this evidence concerns elementary single neuron processes or, alternatively, neuronal ensembles^13–16^ and is insufficient to explain how the granular layer microcircuit units would transform spatially distributed input patterns. A main hypothesis is that the cerebellum input stage operates as a spatially organized *adaptive filter* ^17–20^ implying that plasticity should be able to modify the processing of input patterns in the local microcircuit. Addressing this question requires simultaneous recordings of activity from multiple cerebellar neurons when a bundle of mossy fibers is activated. Under these conditions, short stimulus bursts in the mossy fibers can be used to imitate those that naturally occur during punctuate facial stimulation^21,22^ and generate response bursts in granule cells. Optical fluorescence calcium imaging techniques have recently been used to monitor multiple neuronal activities^23,24^. Here we have used a two-photon confocal microscope based on spatial light modulation (SLM-2PM) that proved capable of recording activity from several granular layer neurons simultaneously and was stable enough to monitor long-term changes in synaptic transmission^25^. Therefore, in principle, SLM-2PM should allow to map spatially distributed granular layer activity with single cell resolution before and after the induction of long-term synaptic plasticity with high-frequency mossy fiber stimuli^26–29^. A further critical step is to extract the single spike patterns of neurons, which cannot be directly measured at the time resolution of calcium imaging but can be inferred using realistic modeling techniques. In recent years, modeling of the granular layer has advanced enough to guarantee a good predictability of local microcircuit dynamics^30–32^. Techniques are therefore mature, in principle, to evaluate the cellular determinates of the adaptive filtering properties predicted by theory in computational units of the cerebellum granular layer.

## RESULTS

### Multi-neuron responses of the granular layer following mossy fiber stimulation

Calcium imaging (with Fura-2 AM) was performed using a spatial light modulator two-photon microscope (SLM-2PM) ^25^ in order to gain insight into the cellular organization of granular layer responses to MF bundle stimulation in acute cerebellar slices. We recorded up to 200 granule cells simultaneously, many of which (normally <=100) responded with fluorescence changes following delivery of short stimulus bursts (10 pulses @ 50 Hz) (Fig.1 A-C). Recordings were carried out both on the sagittal and coronal plane, in order to account for potential asymmetries in granular layer responses. The multi-neuron maps (Δ*F*/*F*_0_ of granule cell responses to MF stimulation) covered a surface that was irregularly rounded both in sagittal and coronal slices and showed activity decreasing from core to periphery. After applying the GABA-A receptor blocker, 10 μM gabazine, the responses increased indicating that Golgi cell synaptic inhibition was limiting the extension and intensity of granule cell activation.

**Fig. 1.**
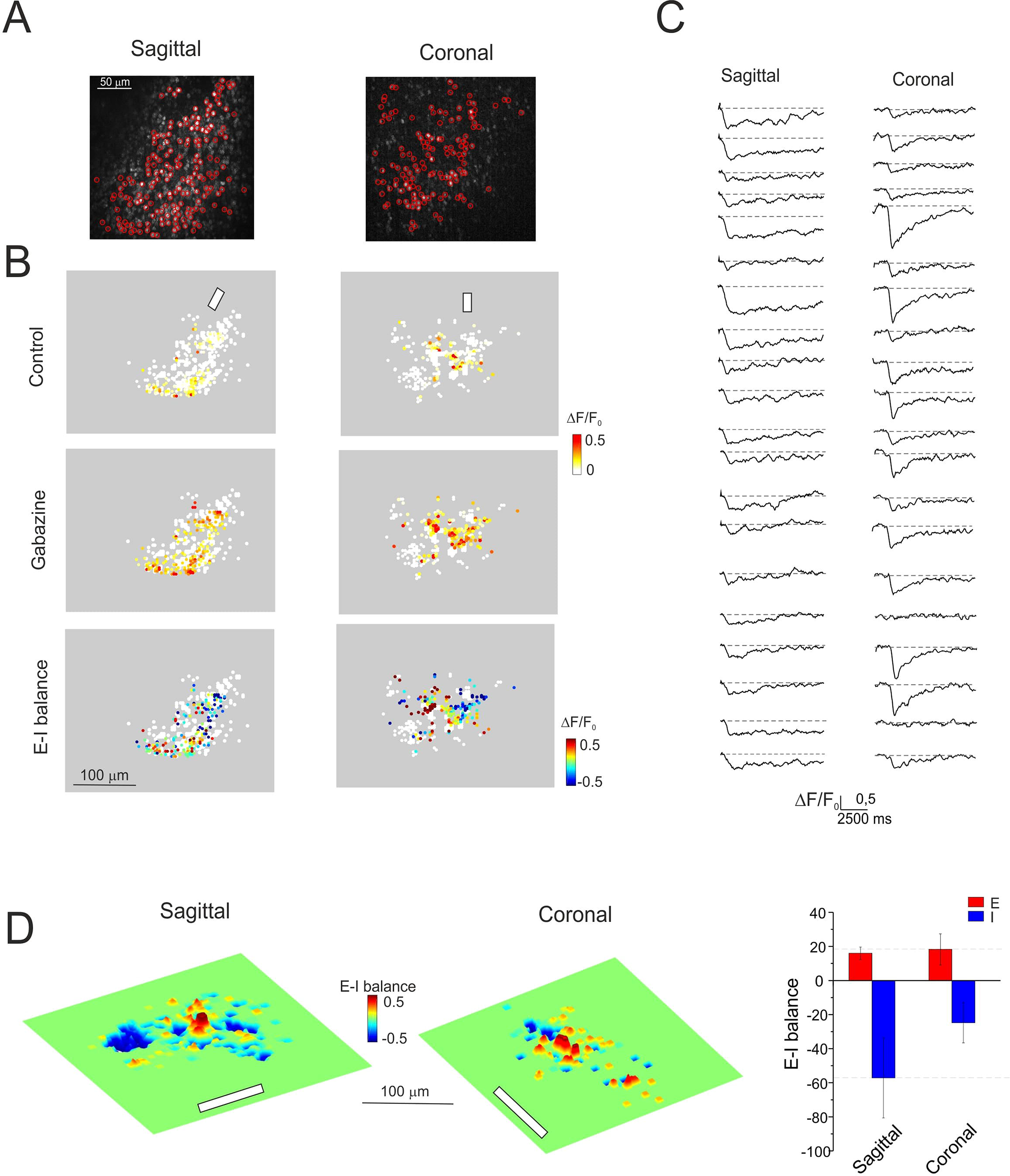
Multi-neuron maps of granule cell activity. **A.** SLM-2PM images of sagittal and coronal cerebellar slices (20x objective, Fura-2 AM bulk loading). Some cells (red circles) are selected for calcium imaging. **B.** Activity maps, taken from the cells selected in A, show on a color scale the peak intensity (Δ*F*/*F*_0_) of the granule cell responses to mossy fibers stimulation (10 pulses@50 Hz, white bars). Each dot corresponds to a neuron. Both in sagittal and coronal slices, the maps show variable intensity of responses from cell to cell and a remarkable increase of activity after applying gabazine (10 μM, continuous bath perfusion). The E-I balance maps (see Methods) show that excitation or inhibition tend to prevail in distinct areas. **C.** Examples of stimulus-induced Ca^2+^ signals simultaneously detected from the soma of different granule cells. Fura-2 AM fluorescence signals appear as ΔF/F_0_ reductions induced by intracellular calcium increase (6 points adjacent-averaging smoothing). **D.** Cumulative E-I balance maps show the spatial profile of granule cells activity (5 sagittal and 4 coronal slices oriented with respect to the mossy fiber bundle, white bars). It should be noted that the most excited areas lye in the core and are flanked by inhibition. The histograms on the right show that the cumulative activations of the E and I areas are similar in sagittal and coronal slices (p=0.8 and p=0.3, respectively; unpaired *t*-test).

The comparison of responses before and after GABA-A receptor blockade allowed to calculate the E-I balance (Fig. 1B). The multi-neuron maps revealed evident core excitation (E-I>0) and widespread peripheral inhibition (E-I<0), confirming initial observations^25^. This arrangement was evident both in the sagittal and coronal planes.

Since multi-neuron maps were rather irregular, the spatial organization of neuronal activity was reconstructed by generating cumulative response maps from several recordings (5 sagittal slices and 4 coronal slices) (see Methods) (Fig. 1D). These cumulative maps closely resembled the center/surround (C/S) organization observed using local field potential recordings and voltage-sensitive dye imaging^13,16^. The global activities (summing up the contribution of all active cells) in regions with either E-I>0 or E-I<0, were similar when comparing sagittal and coronal slices (p=0.8 and p=0.3, respectively; unpaired *t*-test; Fig. 1C). All subsequent experiments were carried out on sagittal slices.

### Spatial organization of long-term synaptic plasticity in the granular layer

Since SLM-2PM recordings proved stable over time^25^, they were exploited to investigate the spatial organization of plastic changes induced by high-frequency stimulation (HFS) of the mossy fiber bundle. The FURA-2 signals were monitored in sagittal slices over a recording time of 60 min, reproducing a protocol similar to that used successfully in whole-cell recordings from granule cells^26–28,33^.

In a first set of experiments (n = 7; Fig. 2A), SLM-2PM recordings were carried out while keeping inhibition intact (control condition). HFS induced rapid and persistent changes in Δ*F*/*F*_0_ of granule cell responses configuring either CaR-P (*calcium-related potentiation*) (99.8 ± 6.5%, n = 26 cells; p<0.01, unpaired *t*-test) or CaR-D (*calcium-related depression*) (−35.1 ± 3.9%, n = 67 cells; p<0.01, unpaired *t*-test). Some granule cells showed changes in Δ*F*/*F*0 <= ± 20% (−3.7 ± 1.6%, n = 47 cells; p=0.1, N.S.) falling within spontaneous response variability (cf. Fig. 2 in ^25^) and were not counted as either CaR-P or CaR-D.

**Fig. 2.**
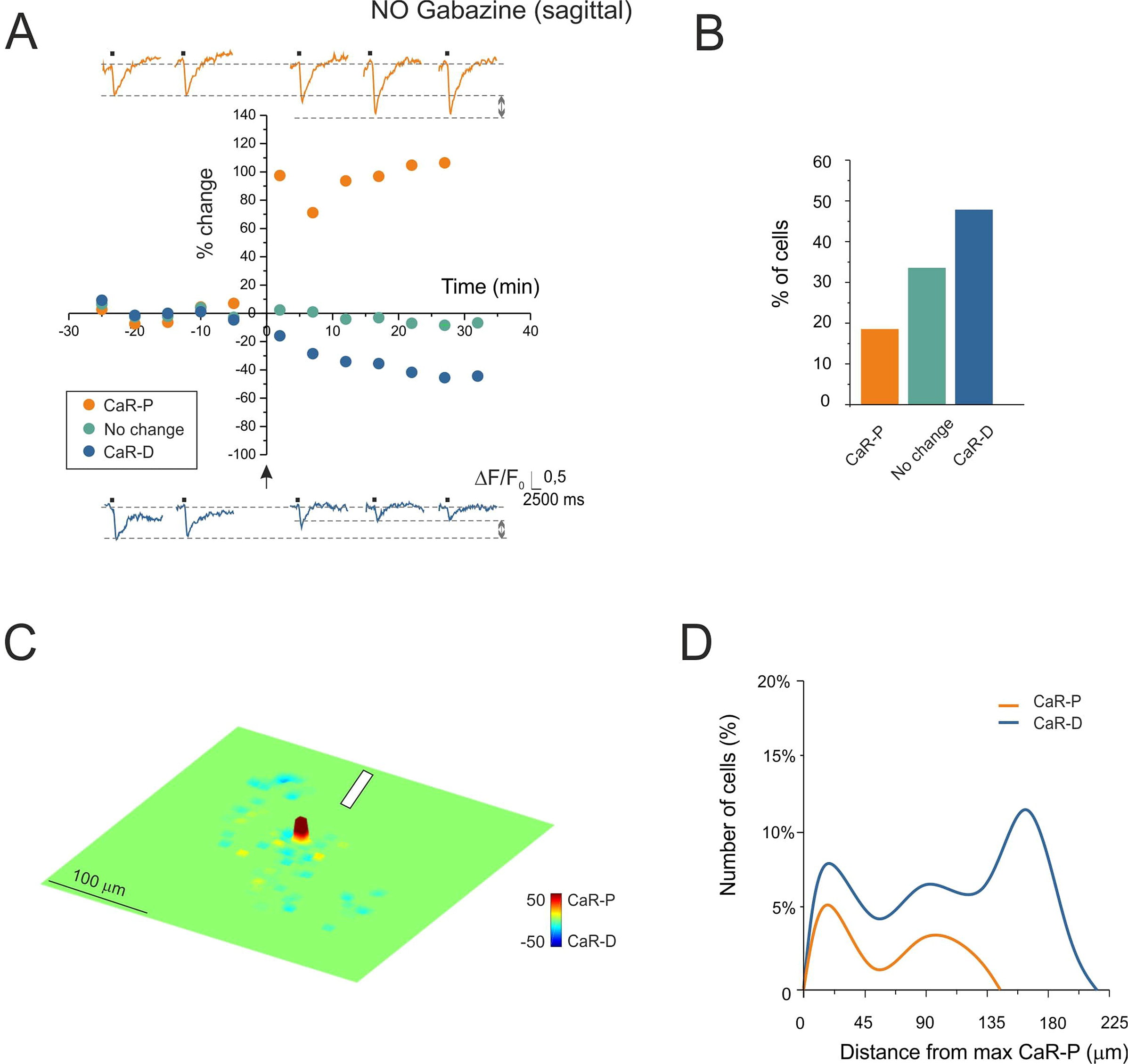
Long-term synaptic plasticity in multi-neuron recordings: control. **A.** Time course of granule cells calcium signal amplitude before and after plasticity induction (HFS was delivered to the mossy fiber bundle at the arrow). Persistent granule cell response variations appear as CaR-P (>20%) or CaR-D (<-20%). The changes <±20% are shown separately. The data points are the average of all the cells falling in the same category taken from available experiments (7 slices, 140 cells; standard errors are small and fall inside the dots). The exemplar traces show Ca^2+^ signals before and after plasticity induction. **B.** Histograms showing the percentage of cells in each category shown in A. **C.** The cumulative plasticity map shows a CaR-P area surrounded by CaR-D (7 slices oriented with respect to the mossy fiber bundle, white bar). **D.** Radial profiles of CaR-P and CaR-D. Note that CaR-D prevails over CaR-P by cell number and extension.

In a second set of experiments (n = 5; Fig. 3A), recordings were carried out with inhibition blocked by 10 μM gabazine. As in control, HFS induced rapid and persistent changes in Δ*F*/*F*_0_ of granule cell responses configuring either CaR-P (90.6 ± 6.1%, n = 217 cells; p<0.01, unpaired *t*-test) or CaR-D (−49.1 ± 2.4%, n = 51 cells; p<0.01, unpaired *t*-test). The cells showing changes in Δ*F*/*F*0 <= ± 20% (−3.6 ± 1.6%, n = 108 cells; p=0.08, N.S.) were not counted as either CaR-P or CaR-D.

**Fig. 3.**
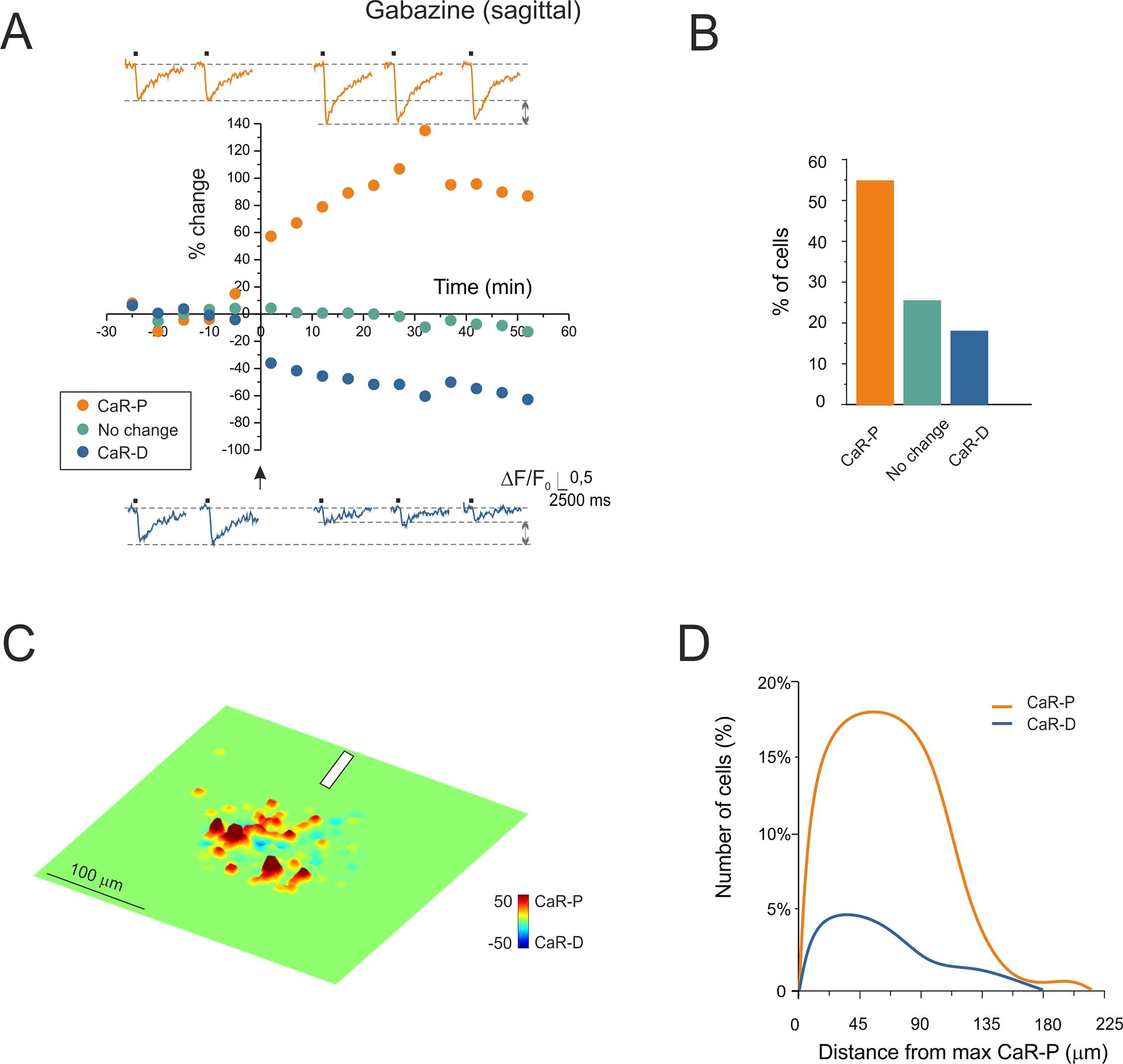
Long-term synaptic plasticity in multi-neuron recordings: gabazine. **A.** Time course of granule cells calcium signals amplitude before and after plasticity induction (HFS protocol delivered to the mossy fiber bundle, arrow). Persistent granule cells response variations appeared as CaR-P (>20%) or CaR-D (<−20%). The changes < ±20% are shown separately. The data points are the average of all the cells falling in the same category taken from available experiments (5 slices, 370 cells; standard errors are small and fall inside the dots). The exemplar traces show Ca^2+^ signals before and after plasticity induction. **B.** Histograms showing the percentage of cells in each category shown in A. **C.** The cumulative plasticity map shows a wide CaR-P area flanked by CaR-D (5 slices oriented with respect to the mossy fiber bundle, white bar). **D.** Radial profiles of CaR-P and CaR-D. Note that, when inhibition is blocked, CaR-P prevails over CaR-D by cell number and extension.

In synthesis, CaR-P and CaR-D were significantly expressed both in the presence and absence of inhibition. The application of gabazine markedly increase the number of cells showing CaR-P rather than CaR-D and therefore inverted the CaR-P/CaR-D ratio (cf. Fig. 2B to Fig. 3B).

In order to visualize the spatial organization of long-term response changes, cumulative plasticity maps were computed from several experiments (see Methods). When synaptic inhibition was intact, the map showed a sharp CaR-P area surrounded by CaR-D (Fig. 2C). When synaptic inhibition was blocked, CaR-P and CaR-D areas largely overlapped (Fig. 3C). These properties reflected in the radial profiles of the number of cells showing CaR-P and CaR-D (Fig. 2D and 3D), that clearly demonstrated the inversion of the CaR-P/CaR-D ratio along with the extension of CaR-P beyond CaR-D when inhibition was blocked.

### Simulation of multi-neuron responses in a model of the cerebellar granular layer microcircuit

Although SLM-2PM calcium imaging provided a unique description of multi-cellular activity and plastic changes organized in space, the data were collected on a plane and single neuron spikes could not be resolved. In order to predict the 3D structure of plasticity with spike-time resolution we used a detailed data-driven model of the cerebellum granular layer (see Methods and Supplemental Material, Fig. s1).

The model was constructed using realistic connectivity and was endowed with detailed neuronal and synaptic properties ^11,29,32,34^. The model was first used to determine the *microcircuit unit* generating responses compatible with experimental recordings *in vitro* using bursts delivered to the MF bundle (10 pulses @ 50 Hz). Similar bursts patterns were shown to occur during punctuate sensory stimulation *in vivo* ^21,35,36,15,22,37^ (Fig. 4A). Simulations were run by activating an increasing number of neighboring glomeruli (see Methods). A C/S structure clearly emerged with >10 glomeruli and its size increased with the number of glomeruli (Fig. 4A). Under the assumption that there is a direct relationship between Δ*F*/*F*0, membrane depolarization and the number of spikes (see Fig. 4 in Ref. ^25^), we compared SLM-2PM recordings to model output. According to simulations, a close matching with the SLM-2PM imaging data of Fig. 1 was obtained when about 50 contiguous glomeruli were activated. The simulations with 50 glomeruli revealed the spatial distribution of spiking activity in the multi-neuron unit (Fig. 4B). Interestingly, with inhibition ON, granule cells showed a variety of firing patterns, with different fists-spike delays and number of spikes, which were clearly correlated with their position in the unit. Conversely, with inhibition OFF, the discharge pattern became more stereotyped, with short delays and high discharge frequencies. The global activity of regions with either E-I>0 or E-I<0 were comparable in sagittal and coronal slices (p=0.75 and p=0.57, respectively; unpaired *t*-test) ruling out asymmetry in simulated as well as in experimental C/S. All subsequent analysis of model simulations was carried out on the whole 3D response unit (see also Supplemental Material, Figs s2-s3).

**Fig. 4.**
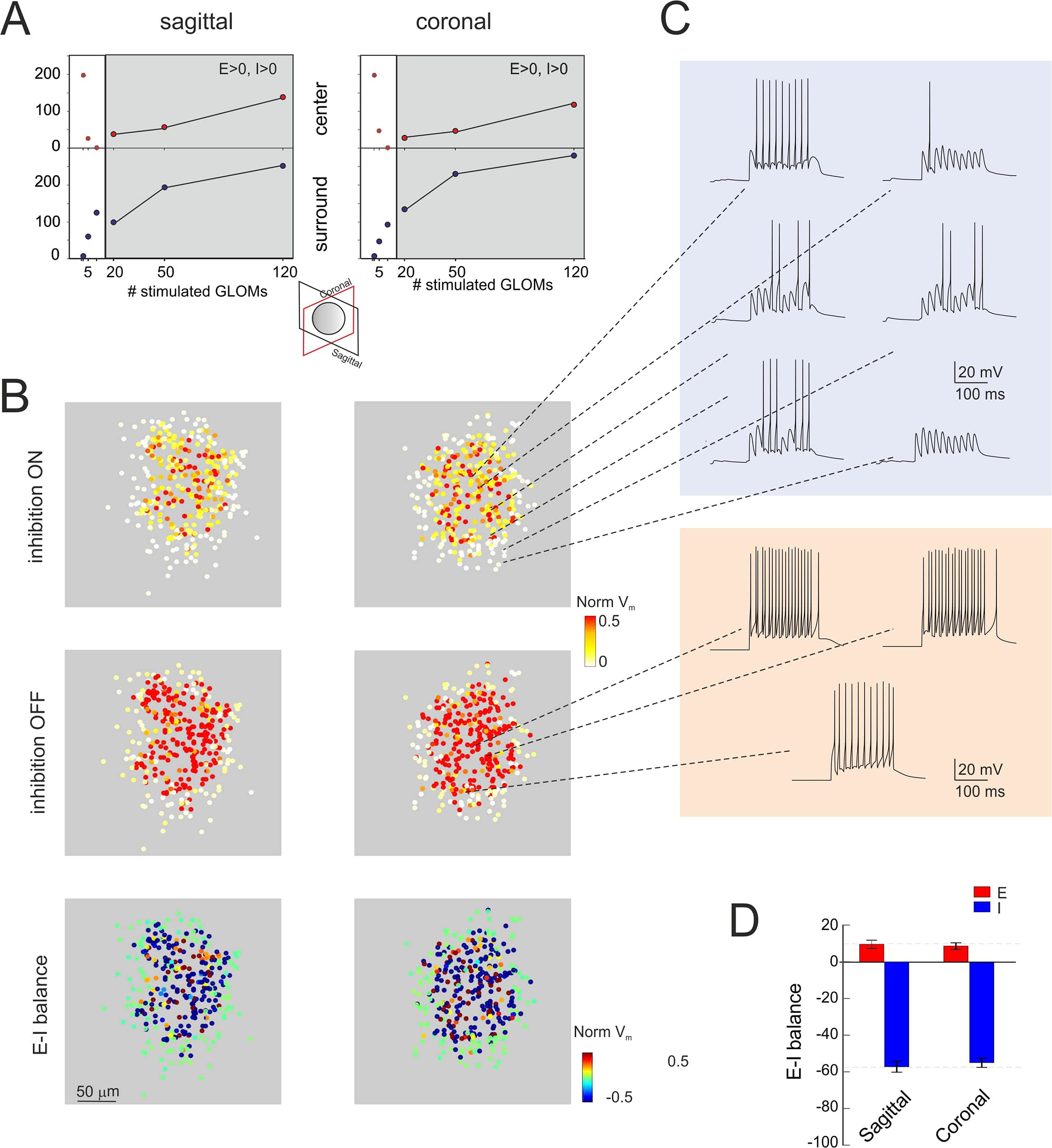
Simulated microcircuit response units. **A.** A realistic data-driven model of the cerebellum granular layer (Methods and Supplemental Material) is used to determine the size of the microcircuit units generating responses compatible with SLM-2PM recordings like those reported in Fig. 1. A typical C/S response structure emerged by activating more than 10 neighboring glomeruli, as shown by the activity maps. **B.** Sagittal and coronal sections of the response unit show the normalized membrane potential (V_m_) of neurons in responses to the activation of 50 contiguous glomeruli (10 pulses@50 Hz). Each dot corresponds to a granule cell of the unit. Both in sagittal and coronal sections, the maps show variable intensity of responses from cell to cell and a remarkable increase of activity after turning inhibition off. The E-I balance maps show distinct areas of prevailing excitation or inhibition. **C.** Examples of simulated voltage traces related to granule cells from different positions in the microcircuit unit. Note that, when inhibition is on, granule cells show a variety of firing patterns, while with inhibition off the pattern is more stereotyped. **D.** The histograms show that the cumulative activations of the E and I areas are similar in sagittal and coronal sections of the 3D activity maps (p=0.75 and p=0.57, respectively; unpaired *t*-test).

### Simulation of plastic changes

One unresolved issue in experimental studies is how synaptic plasticity (long-term potentiation and long-term depression, LTP and LTD) is spatially distributed throughout the microcircuits. The model was therefore used to predict the expression of plasticity based on its cellular mechanisms. In granule cells receiving mossy fiber bursts, the membrane potential change is known to bring about corresponding modifications in Ca^2+^ influx through NMDA receptor-channels, which effectively trigger plasticity induction at mossy fiber - granule cell synapses ^26,38^. LTP and LTD were simulated by measuring the average membrane potential of each granule cell during HFS, calculating neurotransmission changes through the Lisman / Shouval rule^39,40^, and then modifying MF release probability accordingly. This last step accounts for the release probability values estimated by whole-cell recordings and quantal theory calculations^26,41^ (Fig. 5) (see also Supplemental Material, Figs s4-s5).

**Fig. 5.**
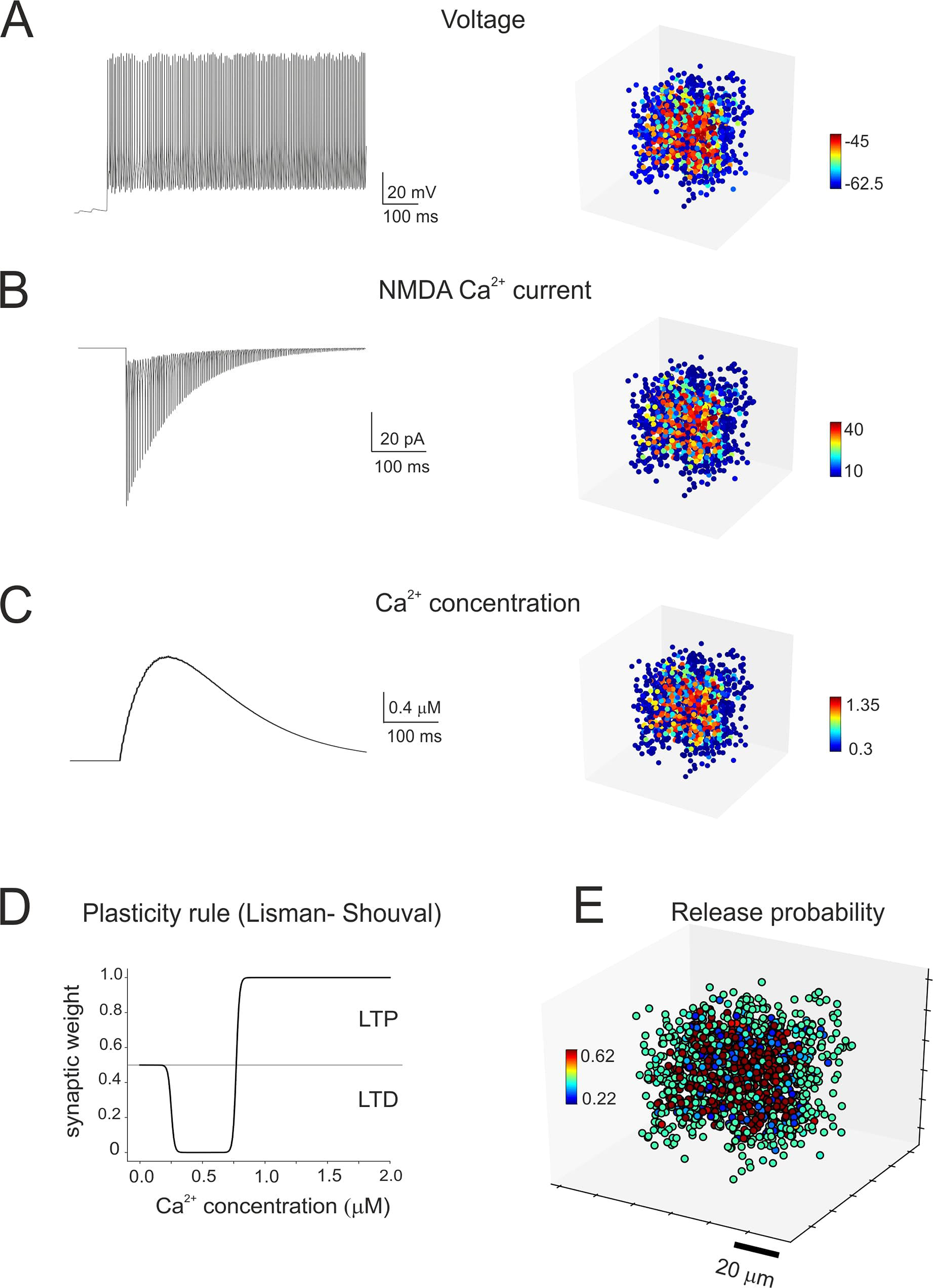
Simulation of synaptic plasticity. **A-C.** Cellular firing caused by HFS (same as in the experimental induction protocols) determines Ca^2+^ influx through synaptic NMDA receptor-channels and the consequent increase in intracellular Ca^2+^ concentration. Membrane voltage (average voltage), NMDA current (integral of the current) and Ca^2+^ concentration (peak concentration) vary from cell to cell depending on local connectivity inside the microcircuit unit, as shown in the 3D activity maps on the left. **D.** The Lisman/Shouval rule is used to calculate the neurotransmission changes induced by the Ca^2+^ concentrations reached during HFS. **E.** The mossy fibers release probability is changed accordingly (here plotted over the receiving granule cells) to account for LTP or LTD expression.

The experiments reported in Figs 2 and 3 were simulated with either inhibition ON or OFF (Fig. 6). Similar to experiments, LTP was dominant in the core and LTD in the periphery and, when inhibition was turned off, the LTP/LTD ratio increased and the core area extended toward the unit’s edge. These results reflect the arrangement of membrane voltage, NMDA current and [Ca^2+^]_i_ during induction (cf. Fig. 5). Since membrane depolarization was normally higher in the core than in the periphery, the probability of having LTP was also higher in the core, while the probability of having LTD was higher in the periphery.

**Fig. 6.**
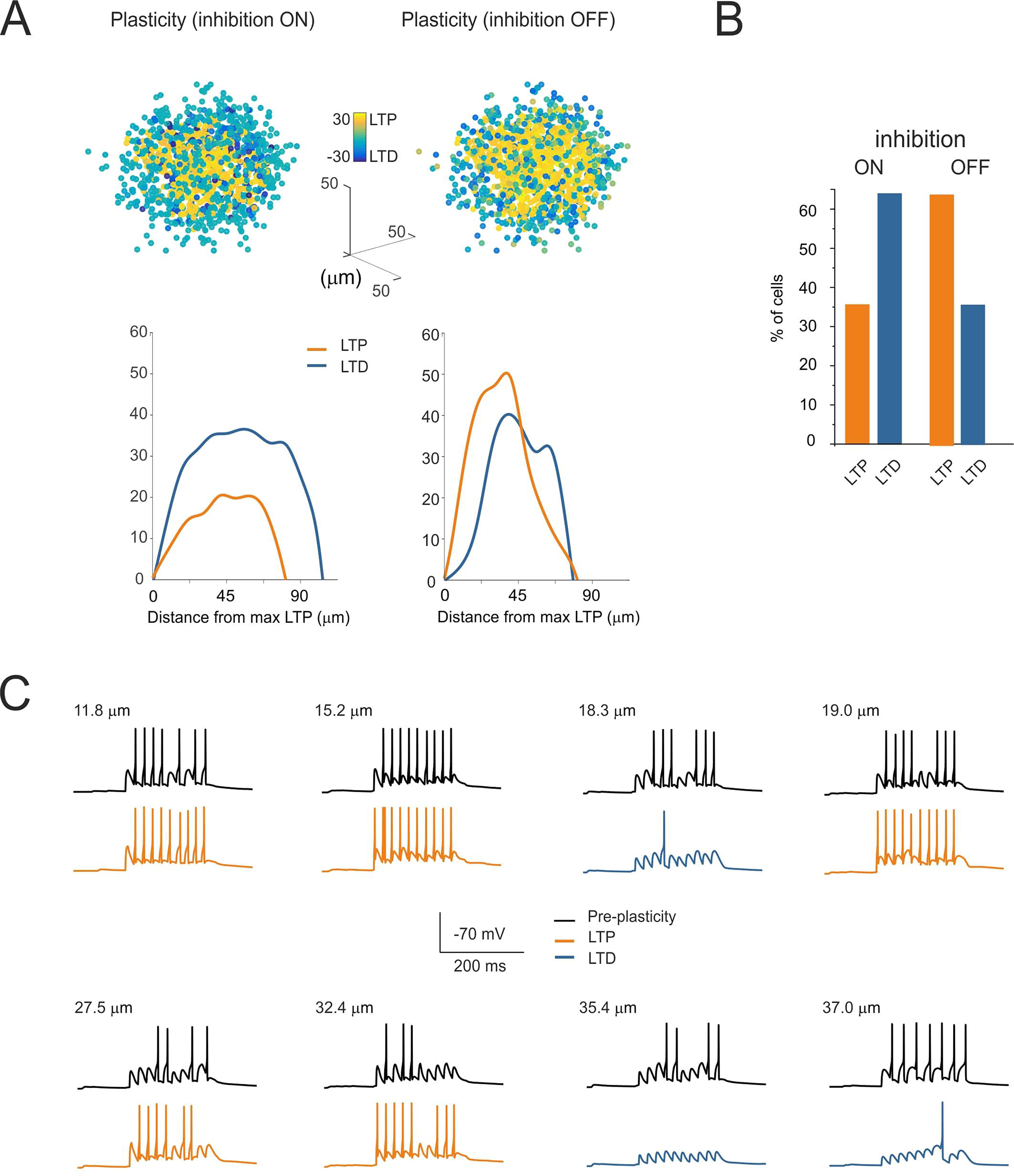
Simulation of plastic changes. **A.** 3D maps of simulated plastic changes with either inhibition on or off. The radial profiles of LTP and LTD are shown at the bottom. Note that, when inhibition is blocked, LTP prevails over LTD by both number of granule cells and extension toward the edge of the recorded area, mirroring experimental observations (cf. Figs 2 and 3). **B.** Histograms showing the percentage of cells in each category shown in A. Note the similarity with experimental observations (cf. Figs 2 and 3). **C.** Exemplar traces of simulated granule cell responses to test stimuli (10 pulses@50 Hz) before and after plasticity (inhibition on). Note that, after plasticity, spike discharges are potentiated and anticipated in the core but are depressed and delayed in the periphery.

A specific advantage of simulations was to get insight into spike timing of the multi-neuron units. Amazingly, while control experiments showed a large distribution of first-spike delays and firing patterns, delay distribution became narrow after plasticity induction and the patterns became more clearly distinguished between those firing strongly and rapidly (mostly in the core) and those firing weakly and slowly (mostly in the periphery). Exemplar traces are shown in Fig. 6C.

### Prediction of spatiotemporal recoding of mossy fiber inputs in the granular layer

The granular layer has been proposed to perform spatiotemporal recoding of the incoming mossy fiber inputs ^8^ but the cellular basis of this operation have remained unclear. Here we have used the model in order to analyze how synaptic plasticity can regulate gain^10^ and lag^11^ of the input-output function. These operations were assessed by applying input bursts (5 spikes) at different frequencies (5-100 Hz) to explore the physiological bandwidth^14^ in control and after the induction of synaptic plasticity.

The response of a unit to different input patterns (fig. 7A) showed that that, after plasticity, the granule cells responded more rapidly, with more spikes and at higher frequency. A quantitative assessment was obtained by measuring the gain (as the number of cells emitting spikes along with spike number and frequency) and the lag (as the time-to-first spike). The multi neuronal unit showed a characteristic behavior (Fig. 7B). When the input frequency was raised, the number of firing cells increased, the gain remained almost constant (variation < 0.1) and the lag decreased considerably (range 20-150 ms). After the induction of plasticity, all parameters changed consistently, leading to a higher number of firing cells, higher gain (~ +0.2) and lower lag (range 10-80 ms). Moreover, after the induction of plasticity, the difference between lower to higher frequencies was more accentuated.

**Fig. 7.**
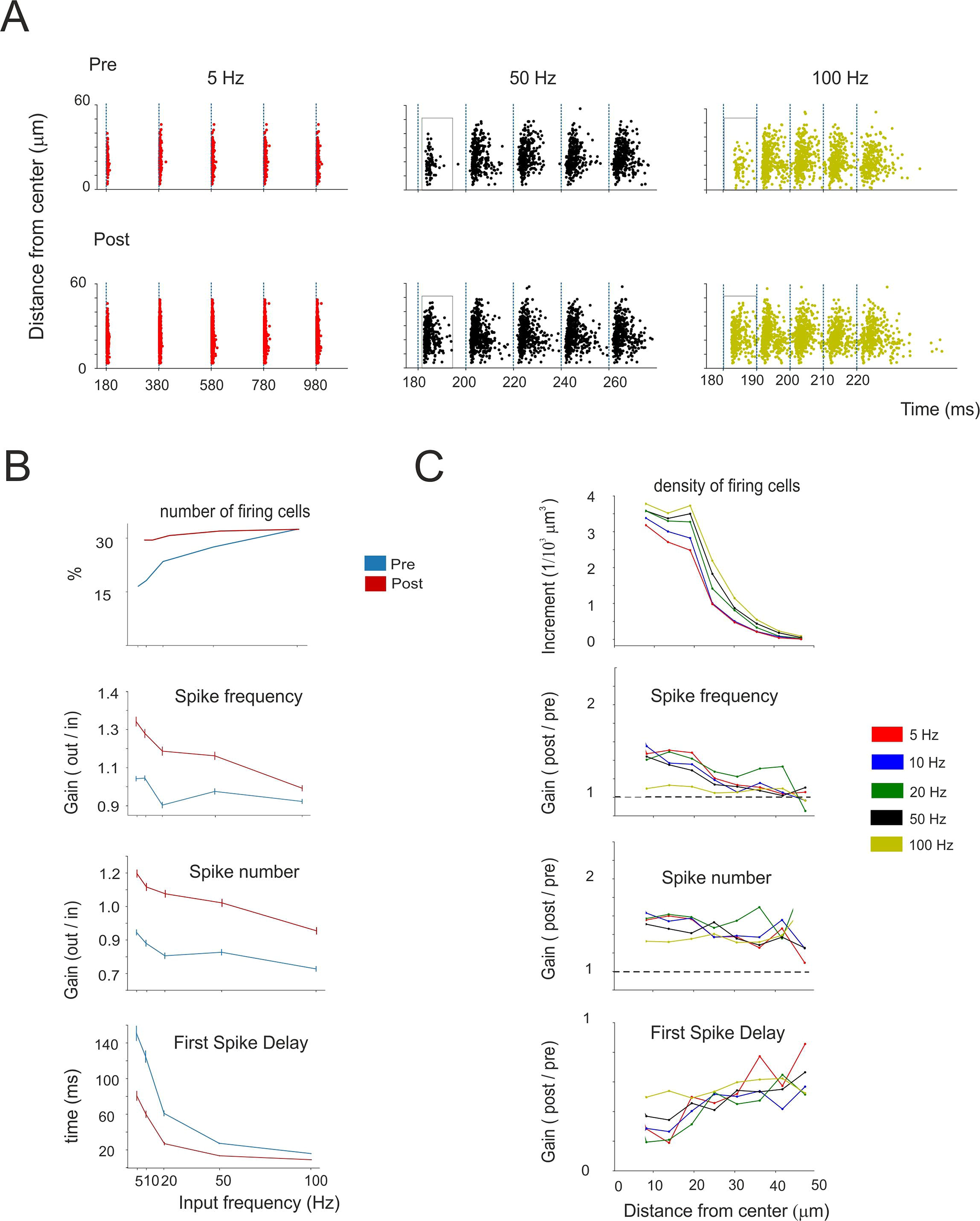
Spike transmission tuning in the response unit. **A.** Raster plots showing the response of all granule cells in a unit to different input patterns (5 pulses@5 Hz; 5 pulses@50 Hz; 5 pulses@100 Hz), both before and after the induction of synaptic plasticity. Note that, after plasticity, the cells respond more rapidly, with more spikes and at higher frequency (boxes highlight these differences for responses to the first input pulse). **B.** Frequency-dependence of the number of firing cells, spike frequency, spike number and first-spike delay, before and after plasticity. **C**. The plots show the increment (post-pre) or the gain (post/pre) after plasticity of the parameters reported in B as a function of distance from the center of the unit. Note that tuning of spike frequency and first-spike delay is more marked in the core of the unit.

At all frequencies, the changes induced by plasticity were more accentuated in the core compared to the periphery of the response unit (Fig. 7C), reflecting the concentration of LTP in the core and of LTD in the periphery. Thus, synaptic plasticity eventually increased the contrast between these two regions of the response unit, with the effect of focusing the transmission channel.

### Simulation of filtering channels tuned by synaptic plasticity

The possibility of having different filtering channels was simulated by comparing three units located in different positions in the network (Fig. 8A). The hypothesis is that, by virtue of the variable wiring with mossy fibers and Golgi cells, the granule cells in these units are in a different “initial state” and will thus develop different membrane potential changes and plasticity (e.g. see ^19,20^). We have therefore stimulated the three units with input spike patterns containing mixed frequency components (e.g. in the theta, alpha, beta and gamma band). The spectral pattern of the firing frequencies transmitted by the three units, before and after plasticity, are shown in Fig. 8B. Clearly the three units are not identical and, after plasticity, unit *I* becomes the best in transmitting at ~50 Hz and ~80 Hz, unit *II* at ~10 Hz, unit *III* at ~70 Hz. Thus, the three units clearly differentiate their transmission properties after plasticity and behave as differential filters.

**Fig. 8.**
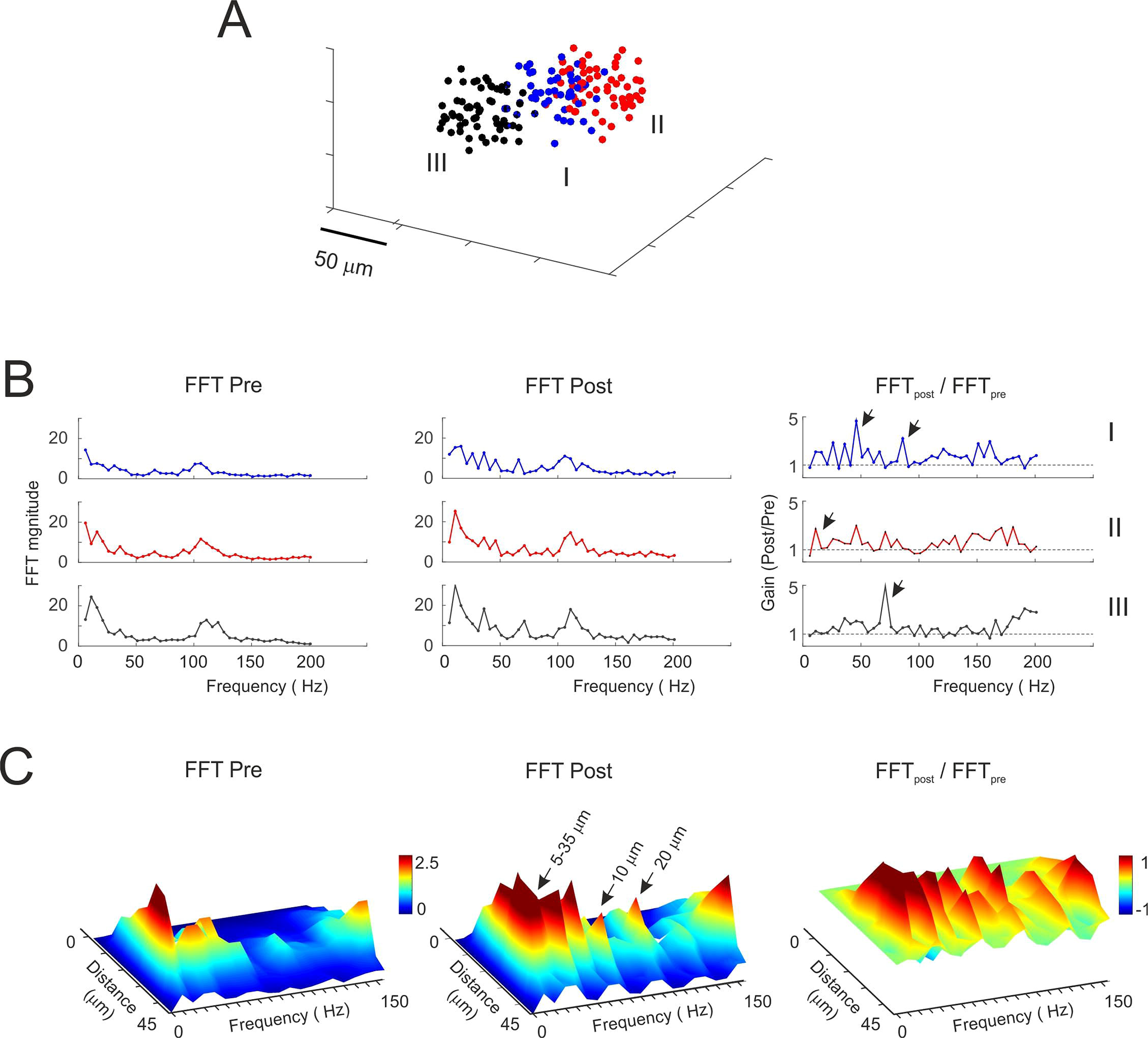
The basis of differential filtering in granular layer response units. **A.** 3D activity maps showing the mossy fiber glomeruli activating 3 units with a mossy fiber pattern containing mixed frequencies (25%@20Hz, 25%@30Hz, 50%@4Hz with superimposed 100Hz bursts). **B**. The plots show the Fast-Fourier Transform (FFT) of the 3 units (all output spikes pooled together) before and after plasticity. The ratio (FFT_post_ / FFT_pre_), for each unit, shows the emergence of preferred frequencies, at which the units have become better in transmitting spikes, e.g. unit #1 at 50 Hz and 80 Hz, unit #2 at 10Hz, unit #3 at 70 Hz (arrows). **C**. The plots show the FFT of unit #1 after separating the output spikes depending on their distance from the center of the unit, before and after plasticity. The ratio (FFT_post_ / FFT_pre_) shows the regions that, for each frequency, change their transmission properties. Note a general improvement of transmission after plasticity with the 40 Hz peak located at 10 μm and the 80 Hz peak located at 20 μm from the center of the response unit.

We then considered how the transmission properties were distributed into the response unit by running the FFT in concentric spherical crowns. (Fig. 8C). Actually, frequencies were differentially filtered at specific distances from the center of the unit and after plasticity, a generalized enhancement of transmission spread over the whole unit.

## DISCUSSION

The spatial filtering properties of the cerebellar granular layer revealed in this work support a critical prediction of the Motor Learning Theory ^8,9,19,20,42^ (MLT). The theory suggests that inputs entering the cerebellum undergo a process of *combinatorial expansion recoding*, in which the signals are separated into different components and transformed before being conveyed to the molecular layer. *Combinatorial expansion* was thought to descend from the divergence/convergence ratio of the mossy fiber - granule cell synapses and *recoding* from the inhibitory control that Golgi cells exert on granule cell activity. Then, adaptation needed for learning was thought to occur through plasticity at the parallel fiber - Purkinje cell synapses. This view has been implemented into the Adaptive Filter Model (AFM^17–20^), in which information is actually split through different channels processing the input signals in the granular layer. Our observations show that such channels can indeed be generated through the contribution of granule cells and Golgi cells aggregated into multi-neuronal units. Interestingly, these channels proved themselves tunable by synaptic plasticity adding a further dimension to the cerebellum granular layer filtering properties.

### The granular layer response units at single cell resolution

When a mossy fiber bundle was activated by input bursts imitating those reported in vivo following punctuate sensory stimulation^21,22^, the granular layer responses were organized in *units* similar to those observed previously using multielectrode arrays ^13^, voltage-sensitive dye imaging ^16^ and two-photon microscopy ^25^. The units showed an irregular rounded shape both in sagittal and coronal sections. The core of the units was more excitable than the periphery and excitation became more prominent and extended when inhibition was blocked pharmacologically. The units underwent remodeling following high-frequency mossy fiber stimulation and this made sharper the difference between core and periphery. The underlying cellular changes were called CaR-P and CaR-D and had magnitude and time course resembling those of LTP and LTD reported at the mossy fiber - granule cells synapse ^26–28,33^. Interestingly, about 20% of the granule cells underwent CaR-P and 50% CaR-D, but this proportion was almost inverted when synaptic inhibition was blocked by gabazine. Moreover, while normally CaR-P involved less cells than CaR-D and was confined to the internal part of the unit, in the absence of inhibition the cells showing CaR-P overtook in number those showing CaR-D and extended till the edge of the unit.

Model simulations at single cell spikes resolution showed that units, similar to those observed with SLM-2PM, could be generated by activating about 50 adjacent glomeruli contacting 300-400 granule cells. These units occupied an irregular rounded tridimensional volume with the most excited cells in the core. The switch-off of Golgi cell inhibition changed the response in a way consistent with the calcium imaging experiments using pharmacological inhibition (cf. Fig. 4 to Fig. 1). The size and properties of these units resemble those obtained from reconvolution of local-field potentials elicited by punctuate facial stimulation^15^, which predicted 11% discharging granule cells at rest and an increase to about 30% during plasticity over about 250 granule cells (cf. Fig. 7B). Therefore, these units obtained *in vitro* and *in computo* are likely to represent the counterpart of the *dense clusters* activated in the granular layer *in vivo*, further supporting their physiological relevance.

### The organization of long-term synaptic plasticity inside the response units

High-frequency stimulation of mossy fibers is known to induce NMDA-receptor-dependent LTP and LTD in granule cells^26^, which was previously shown to conform to the Lisman rule for Ca^2+^-dependent induction^26,38^. The Lisman-Shouval^39,40^ plasticity model showed that the differential distribution of membrane potential, NMDA receptor activation and calcium influx in the granule cells of the units could indeed account for the plastic changes obtained with SLM-2PM recordings. Simulations correctly predicted that LTP was more concentrated in the core and LTD in the periphery of the unit. Moreover, the LTP/LTD ratio was inverted and LTP extended toward the edge of the simulated unit when inhibition was switched-off (further details in Supplemental Material, Fig. s5). Therefore, LTP and LTD predicted by the Lisman-Shouval model were sufficient to explain the spatial properties of experimental CaR-P and CaR-D. Additional mechanisms, like non-synaptic plasticity ^15,28^, intervention of mGlu-Rs and Ca^2+^ release from internal stores ^26,38^, and combined pre- and postsynaptic expression mechanisms ^16,27^ could also contribute to change the LTP/LTD landscape in the unit.

### Model prediction of spike filtering properties in granular layer units

Model simulations predicted that granular layer units express filtering properties tuned by long-term synaptic plasticity. Raising mossy fiber frequency from 5 Hz to 100 Hz almost doubled the number of firing granule cells (gain x2), explaining the corresponding increase in VSD signals reported previously ^14^. The corresponding change in spike transmission (frequency and number) was smaller (gain x0.1), since spike discharges were limited by Golgi cell inhibition (cf. Fig. 4). Interestingly, there was a remarkable modulation of first spike delay, which almost halved (lag x2). Thus, the input frequency prominently modulates the number of granule cells and the time to first spike and much less so the spike discharge frequency and number.

The positive effects of plasticity (increased number of firing granule cells, increased spike transmission gain, reduced first spike delay) occurred in the core of the unit, while the opposite occurred in the periphery. This created a spatial filter channeling high-gain rapid information transfer through the core with enhanced signal contrast.

During plasticity, the changes in granule cell responses (increased number of firing granule cells, increased spike transmission gain, reduced first spike delay) were more evident at low than high frequencies. Plasticity generated therefore a low-pass filter, which may engage multiple molecular changes ^43^ and show resonance peaks in the theta band ^30,44^. During plasticity, first spike delay almost halved reflecting presynaptic regulation of neurotransmitter release probability ^11^ (cf. Fig. 6).

By virtue of variability in mossy fiber - granule cell - Golgi cell wiring, different units were not identical with respect to their filtering properties and this caused differences that became quite relevant following plasticity. These simulations therefore strongly support the concept that mossy fiber - granule cell plasticity is fundamental to tune the filtering properties of the cerebellar granular layer. The recently discovered granule cell subtypes (adapting, non-adapting and accelerating) may further bias the unit toward high-pass or low-pass filtering depending on the expression of TRPM4 channels ^45^.

### Conclusions

The microscopic analysis of granular layer transmission properties modulated by long-term synaptic plasticity supports the *adaptive filter model* of the cerebellum ^19,20,46^, although with differences from original believes. First, granular layer filtering involves spatially organized multicellular units including 50 glomeruli and 300-400 active granule cells. Secondly, granular layer filtering is tuned by local mechanisms of long-term synaptic plasticity. Thirdly, this plasticity involves both LTP and LTD in a spatially organized manner. These concepts support the hypothesis that the granular layer can form multiple signal filtering channels regulating spike delay and discharge gain^12^. These channels can focus the signal in “vertical beams” travelling toward the molecular layer and thereby activate selective groups of Purkinje cells (see^47–49^). This plasticity-controlled distribution of spike timing and discharge can remarkably impact on mutual information transfer at the cerebellum input stage^50,51^ and on subsequent computations in the molecular layer and Purkinje cells.

## Methods

In this work, experiments and simulations were co-designed so that the experiments were analyzed trough a realistic model of the underlying microcircuit. Results from experiments and simulations were analyzed using similar routines to enhance a direct comparison.

### Experimental recordings

Two-photon calcium imaging recordings from the cerebellar granular layer were performed using an SLM-2PM microscope that allows simultaneous multi-neuron recordings ^25,52^. All experimental protocols were conducted in accordance with international guidelines from the European Union Directive 2010/63/EU on the ethical use of animals and were approved by the ethical committee of Italian Ministry of Health (639.2017-PR; 7/2017-PR).

### Slice preparation and solution

Cerebellar granular layer activity was recorded from the vermis central lobe of acute parasagittal or coronal cerebellar slices (230 μm thick) obtained from 18- to 25-day-old Wistar rats of either sex. Slice preparation was performed as reported previously^25,43,53^. Briefly, rats were decapitated after deep anesthesia with halothane (Sigma, St. Louis, MO), the cerebellum was gently removed and the vermis was isolated, fixed on a vibroslicer’s stage (Leica VT1200S) with cyano-acrylic glue and immersed in cold (2-3°C) oxygenated Krebs solution containing (mM): 120 NaCl, 2 KCl, 2 CaCl_2_, 1.2 MgSO_4_, 1.18 KH_2_PO_4_, 26 NaHCO_3_, and 11 glucose, equilibrated with 95% O_2_–5% CO_2_ (pH 7.4). Slices were allowed to recover at room temperature for at least 40 min before Fura-2 AM staining. For this purpose, a 50 μg aliquot of Fura-2 AM (Molecular Probes, Eugene, OR, USA) was previously dissolved in 48 μl of Dimethyl Sulfoxide (DMSO, Sigma Aldrich) and 2 μl Pluronic F-127 (Molecular Probes) and mixed with 2.5 mL of continuously oxygenated Krebs solution. The slices were placed in this solution and maintained at 35°C for 40 minutes in the dark for bulk loading ^54,55^. Then the slices were gently positioned in the recording chamber and immobilized using a nylon mesh attached to a platinum Ω-wire to improve tissue adhesion and mechanical stability. Oxygenated Krebs solution was perfused during the whole experiment (2 mL/min) and maintained at 32°C with a Peltier feedback device (TC-324B, Warner Instruments, Hamden, CT).

### Multi-neuron two-photon calcium imaging

Two-photon calcium imaging experiments were performed through a spatial light modulator two-photon microscope (SLM-2PM)^25,52^. The SLM-2PM uses a spatial light modulator device (SLM, X10468-07, Hamamatsu, Japan) to perform computer-driven holographic microscopy. This device allows to modulate the phase of a coherent laser-light source (Chameleon Ultra II, Coherent, USA) and to generate different spatial illumination patterns on the sample plane, by taking advantage of optical Fourier transforms. Custom-developed routines written in Python (PSF, 9450 SW Gemini Dr. Beaverton, OR 97008, USA) and based on the iterative-adaptive Gerchberg-Saxon algorithm were used to compute the phase distribution corresponding to a desired illumination pattern^56,57^ (for more details see ^52^). In this way it is possible to split the laser beam into multiple beamlets that can be simultaneously directed onto different points of interest in the sample, thus allowing to record from multiple neuron simultaneously, while maintaining the single cell resolution.

### Experimental data acquisition and analysis

Experiments were performed at depths of ~50-60 *μ*m, using a 20X 1.0 NA water-immersion objective (Zeiss Plan-APOCHROMAT) while the laser power was set ≤ 6 mW/laser beamlet.

Two-photon images were acquired through a high-spatial resolution CCD camera (CoolSnap HQ, Photometrics, Tucson, USA) that covered a field of view of about 220×300 *μ*m^2^; SLM-2PM acquired images were used to identify granule cells somata positions and to select cells for the subsequent calcium signals acquisition^25,52^. Granule cells activity was elicited by electrical stimulation of the mossy fiber bundle (10 pulses@50 Hz repeated 3 times at 0.08 Hz to improve the signal-to-noise ratio, S/N) with a large-tip patch-pipette (~10-20 *μ*m tip) filled with extracellular Krebs’ solution, via a stimulus isolation unit. The stimulus-induced fluorescence calcium signals were acquired through a high-temporal resolution CMOS camera (MICAM Ultima. Scimedia, Japan), connected through an I/O interface (BrainVision, Scimedia, Japan) to a PC controlling illumination, stimulation and data acquisition. The final pixel size was 4.6 × 4.6 *μ*m. Data were acquired at a frequency of 20 Hz and displayed by Brainvision software. Granule cells showed calcium signal variations peaking around 150 ms after the stimulus ^25,52^. The stimulus-induced fluorescence signals were analyzed off-line using custom-written software in Matlab (MathWorks, Natick, MA) by evaluating the peak amplitude fluorescence variations normalized to *F*_0_, i.e. (Δ*F*(*n*) */F*_0_=(*F* (*n*) − *F*_0_)/*F*_0_, where *F_0_* was the mean resting fluorescence sampled for 1400 ms before triggering electrical stimulation and *F(n)* was the fluorescence intensity at the *n*th frame. Given a peak amplitude of 0.2-0.9 and a noise standard error of 0.05-0.08, the Δ*F*/*F*_0_ S/N was about 8-fold, ensuring a reliable measurement of peak response amplitude.

In a set of experiments the granule cells calcium responses were acquired before and after the perfusion of a GABA-A receptors inhibition blocker (10 μM gabazine, SR95531, Abcam). In another set of experiments the granule cells calcium responses were acquired before and after the delivery of a high-frequency mossy fibers stimulation protocol (HFS, 100 pulses @ 100 Hz, known to induce long-term synaptic plasticity at the mossy fiber-granule cells synapses 26, 28, 38), either in control (Krebs) or with inhibition blocked (Krebs + gabazine).

The neurons were considered to show long-term changes, when their Δ*F*/*F*_0_ persistently changed by >± 20% after HFS.

The excitatory-inhibitory (E-I) balance maps were computed was computed using custom-written software in Matlab (MathWorks) for each experiment as the difference between E_norm_ and I_norm_ normalized by E_norm_ (E-I = (E_norm_ - I_norm_)/ E_norm_), where E_norm_ is the response intensity in control condition and I_norm_ is the response intensity variation after gabazine perfusion (both normalized to the maximum response), so that E-I values ranged from 1 (maximal excitation) to −1 (maximal inhibition) (cf. ^13^).

Cumulative response maps were computed by centering the individual maps on the cells showing maximum response in control and by aligning them along the mossy fiber bundle. Cumulative plasticity maps (i.e. those obtained by comparing responses amplitude before and after plasticity induction) were computed by centering the individual maps related on the peak value (maximum potentiation) and by aligning them along the mossy fiber bundle. The cumulative maps were smoothed with a sliding box filter (3 × 3 pixels).

Data are reported as mean ± standard error of the mean (S.E.M.) and statistical comparisons are done using paired and unpaired Student’s *t* test, unless otherwise indicated (not significant at p>0.05).^6^

### Granular layer circuit modeling and simulation

Simulations of the cerebellar granular layer activity were performed using a realistic computational model that allows single-cell resolution of neuronal activity. The model, by incorporating new neuronal and synaptic mechanisms, provides a substantial update and extension of our previous model ^30^. The model is written in Python-NEURON (Python 2.7; NEURON 7.3) ^58,59^ and is scalable, so that changes in network size lead automatically to rescaling the number of networks elements and their connectivity. In this way, the model achieves the scale required for simulation of salient operations in the cerebellar network ^32^, including ~400*10^3^ neurons and ~2*10^6^ synapses. Exploratory simulations were performed on a 72 cores/144 threads cluster (six blades with two Intel Xeon X5650 and 24 Gigabyte of DDR3 ram per blade). During simulations, the time step was fixed at 0.025 ms and the NEURON multi-split option was used to distribute computation corresponding to different cells over different cores (http://www.neuron.yale.edu/phpBB/)^58^. Massive simulations were run on the BluGeneQ class supercomputer JUQUEEN at the Julich computational center within the Human Brain Project using resources allocated through the PRACE project “cerebellum”.

The granular layer network was constructed on the basis of detailed anatomical and functional information^60–68^) and using models of neurons and synapses including biophysical representations of membrane ionic channels and receptors^44,11,29,69,70,34,71^. These neuronal and synaptic models have been extensively validated using electrophysiological and imaging data. This allowed to develop a realistic network model, in which the large number of parameters is constrained to biology. In this bottom-up approach, the functional properties of the network emerge from the properties of constitutive elements and from their synaptic organization (e.g. see^72–74^). The availability of input spike patterns and neuronal responses *in vivo* (^21,22,37,75,76,15^ has allowed to simulate granular layer network dynamics under conditions representative of natural activity states. The network responses were validated by comparison with two-photon microscopy recordings ^25^ (this paper) as well as with previous LFP ^15^, MEA ^13^ and VSD ^77^ recordings of network activity.

### General properties and network architecture

While being based on a previous network design^30^, the size of the model was increased significantly for representing a cerebellar slice with a thickness of ~ 150 μm (800 μm × 800 μm × 150 μm), with 384000 granule cells, 914 Golgi cells and 29415 mossy Fibers /Glomeruli. Moreover, the cellular and synaptic mechanisms have been updated. The main extension were the introduction of excitatory connections from granule cells to Golgi cells through the ascending axon (aa) ^78^, of inhibitory connections among Golgi cells ^79^, of gap junctions between Golgi cells ^80,81^, of the orientation of the Golgi cell axon ^62^. The convergence / divergence ratios for each connection type have been update accordingly (see Table s1 in Supplemental Material).

Network connections were constructed using precise rules, yet allowing the number of connections and synaptic weights to show statistical variability (Gaussian distribution: mean = 1, s.d. = 0.4; see ^82^) and no systematic differences were observed using different seeds for parameter randomization (not shown). Background noise in the network was generated only by pacemaking in Golgi cells (see e.g.^21,22,83^), since granule cells are silent at rest in slice preparations. Neurons and synapses were endowed with multiple receptor and ionic channel-based mechanisms, allowing an accurate representation of neuronal firing. The synapses were endowed with neurotransmitter diffusion mechanisms and with a representation of vesicle cycling, generating spillover and developing short-term facilitation and depression. Details on model construction and validation are given in Supplemental Material.

### Model of synaptic plasticity

In the present work, long-term plasticity at the MF-GrCs synapse has been reproduced increasing (LTP) or decreasing (LTD) the release probability (*p*) according to the rules derived from the Lisman - Shouval model (Lisman, 1989; Shouval et al., 2002). Simulations of synaptic plasticity reproduced the applied experimental protocol:

1. Simulation of network activity *before* induction of plasticity; a 10-spikes burst @50Hz was delivered over 50 contiguous glomeruli, Release probability (*p*) was equal to 0.42 for all the MF-GrC synapse
2. Simulation of plasticity-induction protocol; 100spikes @100Hz delivered over the same set of glomeruli. Single-cell voltage traces were recorded and used to estimate plasticity according to the Lisman - Shouval approach (see below).
3. Simulation of network activity *after* induction of plasticity; same stimulation protocol as in 1. For each MF-GrC synapse, *p* changed according to 2.

Details on the synaptic plasticity model are given in Supplemental Material.

### Simulated data analysis

Activity response maps were used to calculate E, I and E-I balance as with the experimental data, but in this case starting from the average cell membrane potential. Several 3D maps were generated using activity related parameters, including membrane voltage, spike discharge frequency, intracellular calcium and presynaptic release probability. The filtering properties were analyzed using a FFT of the output signals Matlab (MathWorks, Natick, MA).

## Acknowledgments

This research was supported by the European Union’s Horizon 2020 Framework Programme for Research and Innovation under Specific Grant Agreement 720270 (Human Brain Project SGA1) and 785907 (Human Brain Project SGA2), with specific involvement of the Neuroinformatics Platform, Brain Simulation Platform, HPAC Platform and the Mouse Data (SP1) and Brain Modelling (SP6) subprojects. The work was also sponsored by the MNL project of the Centro Fermi (Rome, Italy) and the European supercomputing PRACE Project 2018184373.

## Author Contributions

Author contributions: M.T. performed SLM-2PM experiments and analyzed the data; S.C. designed the models and performed the simulations; all authors contributed to paper writing and revision; E.D. coordinated research and wrote the final version of the paper.

