## Supplemental text for "Cellular-resolution mapping uncovers spatial adaptive filtering at the cerebellum input stage"

Supplemental material

***GRANULAR LAYER MODEL CONSTRUCTION AND VALIDATION***

***List of abbreviations***

GrCs, granule cells;

GoCs, Golgi cells;

GLOMs, glomeruli;

AA, ascending axon;

PF, parallel fibers;

MF, mossy fibers.

***Network architecture and connectivity***

The model comprised 383999 GrCs and 915 GoCs, i.e. about 1% of the whole mouse cerebellar cortex. Similar to previous versions ^1^, the one used here included the fundamental connections between MF, GrCs and GoCs and was accurately rewired and parameterized to account for recent experimental observations. These included GrC-GoC connections from both AA and PF synapses ^2^, GoC-GoC inhibitory synapses ^3^ and GoC-GoC gap-junctions ^4, 5^. The GoCs received PFs oriented along the transverse axis. The GoC axonal plexus was drawn as an ellipsoid flattened on the sagittal plane ^6^, where it was connected to GLOMs according to experimentally derived rules ^7^. The connectivity between MFs, GrC dendrites and GoC axons in the glomeruli (GLOMs) and between GrCs and GoC dendrites was reconstructed using appropriate convergence-divergence ratios accurately parameterized on anatomical and physiological data (see Table s1 for details).

The generation of the model network occurred in three steps: (1) calculating the number of constitutive elements, (2) distributing the elements in space, and (3) connecting the elements.

(1) Starting from a GrC density of 4 × 106 /mm^3^, the density of GoCs was calculated to be ∼9000/mm^3^ in order to respect the ratio 1:430 reported by ^8^. The density of GLOMs was calculated from the convergence/divergence ratio of the MF-GrC connections. Each GLOMs includes a mean of 53 dendrites from different GrCs and each GrC emits on average four dendrites ^9^. The density of GLOMs was calculated as (4 × 106 /mm^3^ × 4 dendrites)/(53 dendrites/GLOMs) ≈ 3 × 105 /mm^3^.

(2) After calculating the number of constitutive elements, these were placed into the network volume with coordinates drawn from a uniform random distribution. The innervation territories for GoCs were delimited using a geometrical description of the dendritic and axonal fields with respect to the soma ^6^.

(3) The network connections were generated by applying simple rules, most of which can be directly extracted from original works on cerebellar architecture (e.g. see ^10^). (*a*) The GrC dendrites could not reach GLOMs farther than 40 µm (mean dendritic length 13.6 µm). (*b*) A single GrC was not allowed to project more than one dendrite inside the same GLOMs. (*c*) Only one GoC axon was allowed to enter a GLOMs forming inhibitory synapses on all the afferent GrC dendrites. (*d*) A GoC axon entering into one GLOMs was prevented from accessing the neighboring GLOMs sharing GrCs with the first GLOMs. This prevented a GrC from being inhibited twice through the same GoC, a case that does not seem to hold experimentally (see ^11^). (*e*) Each GoC was allowed to access at most 40 GLOMs resulting in a maximum ∼2000 GrCs inhibited by the same GoC. (*f*) GoCs received excitation from ~65 glomeruli, ~4280 GrCs through PFs and 400GrCs through AAs. (*g*) GoC-GoC inhibitory connections and (*h*) GoC-GoC gap-junctions were organized in the same way: each GoC received from (and projected to) ~145 GoCs. By virtue of their extended dendritic fields, GoCs turned out to be connected beyond the volume occupied by the GrCs reached by the same MFs. Since the probability of getting connected to MFs was higher in the core of the bundle, the density of GrC excited by MFs decreased from core to periphery while the probability of GrC being inhibited remained high also in peripheral areas. It should be noted that the rules d and e are clearly simplications (e.g. see ^12^), whose implications could be further explored in the future.

***Table s1***

|  | Divergence | Convergence | reference |
| --- | --- | --- | --- |
| mfs → Granules | 1 : 51.93 (sd = 3) | 3.97 : 1 (sd = 0.72) | ^61^ |
| mfs → GoCs | 1: 1.55 (sd = 1.28) | 64.99 : 1(sd = 0.04) | ^61^ |
| GoCs → mfs | 1: 32.18 (sd = 10.94) | 1: 1 | ^61^ |
| pfs → GoCs | 1: 9.15 (sd = 3.15) | 4281.99 : 1 (sd = 0.09) | ^78^ |
| aa → GoCs | 1: 0.95 (sd = 0.98) | 400 : 1 | ^78^ |
| GoC→ GoC | 1:145.5 (sd = 36.3) | 145.5 : 1 (sd = 36.3) | ^79^ |
| GoC gap junction | 1:145.5 (sd = 36.3) | 145.5 : 1 (sd = 36.3) | ^80, 81^ |

***Table s1: Divergence and Convergence ratios in the model.*** The convergence and divergence ratios and their standard deviation are reported for each synapses along with the reference paper from which the data have been taken. These data have been used to generate model connectivity.

***Single cell and synaptic models***

The GrC and GoC models derived from previous models, which had been carefully tested against available experimental results in slices ^13-16^. These models were able to reproduce all the details of spike shape, timing and frequency in response to current injection and synaptic stimulation. The synaptic models were adapted from the original scheme reported by ^14^ and were able to reproduce the kinetics and size of the EPSCs and IPSCs during repetitive synaptic transmission at the different synapses. These models accounted for vesicular dynamics, neurotransmitter spillover and receptor gating (including multiple closed, desensitized and open states) but not for quantal release mechanisms. The dynamics of synaptic responses were fully determined by the kinetic constants of synaptic and neuronal models. Given the short distances travelled by the spikes, axonal conduction times were considered negligible. Transmission delay was 1 ms for all the synapses.

In order to conform to in vivo conditions, all models had to be adapted from their original temperature T_orig_ to T_sim_ = 37°C using the correction factor Q10=(T_sim_–T_orig_)/10 (^17^; see also ^18-20^). We have used: Q10 = 3 for ionic channel gating, Q10 = 2.4 for receptor gating, Q10 = 1.5 for ionic channel permeation, Q10 = 1.3 for neurotransmitter diffusion, Q10 = 3 for Ca^2+^ pumps and buffers, Q10 = 1.3 (GrC) or 1.7 (GrC) for intracellular Ca^2+^ diffusion. Following adaptation at 37°C, the models were in matching with recordings at this same temperature (data not shown).

The GrC model was adapted from ^14^ by applying appropriate Q10 corrections. In addition, the GABA leakage conductance was increased by two times (60 µS/cm^2^), the inward rectifier K^+^ conductance was increase by 1.5 times (1350 µS/cm^2^) and the leakage reversal potential was adjusted to restoring resting potential to −70 mV (see ^13^). With this asset, the GrC model properly reproduced responses to current injection at 37°C (data not shown) and spike trains observed in vivo ^21, 22^.

The GoC model was adapted from ^15, 16^ by applying appropriate Q10 corrections. Without needing any further change, the GoC model properly reproduced responses to peripheral stimulation observed in vivo ^23^.

The MF-GrC synapses take part to the formation of the cerebellar GLOMs and are glutamatergic and activate AMPA and NMDA receptors. The release, diffusion and ionic receptor mechanisms were the same reported by ^14^. Using a probability of release of 0.6, the model was able to faithfully reproduce postsynaptic currents recorded at 37°C in vitro ^24^ and in vivo ^21, 25^. The time constant of the recovery from depression, τ_REC_ = 8 ms, was derived from in vivo measurements ^22^ and allowed to reproduce natural dynamics of short-term plasticity (the time constants of presynaptic facilitation and vesicle inactivation were set to τ_facil_ = 5 ms and τ_I_ = 1 ms, respectively).

The MF-GoC synapses are similar in several respects to the MF-GrC synapses. They are also located within the cerebellar GLOMs ^10^ and are glutamatergic activating both AMPA and NMDA receptors ^26^ ^2^. The MF-GoC synapse was adapted from the MF-GrC synapse model (see above) to reproduce a peak postsynaptic current of −66 pA ^2^. Release probability and vesicle cycling parameters were set at the same values as at the MF-GrC synapse.

The GrC-GoC synapses are formed by PFs onto GoC apical dendrites in the molecular layer ^27^. These glutamatergic synapses activate AMPA, NMDA and kainate receptors ^28-30^. During repetitive stimulation, the AMPA current shows synaptic depression while the kainate and NMDA currents show slow temporal summation. AMPA and NMDA currents were taken from the MF-GrC synapses and the kainate receptor current was modified from the AMPA kinetic scheme. Release probability was 0.1 and vesicle cycling parameters were set at the same values as at the MF-GrC synapse. The AA contacts GoC basolateral dendrites in the granular layer ^2^; these synapses activate AMPA and NMDA only; their maximal conductance was estimated to be ~2 times higher than AMPA and NMDA currents of PF-GoC synapses. Also in this case, AMPA and NMDA currents were taken from the MF-GrC synapse; release probability and vesicle cycling were set at the same values, too.

The GoC-GrC synapses are GABAergic and impinge on GrC dendrites within the GLOMs. GABAergic neurotransmission was modeled based on ^31^. The GABA-A receptor schemes comprised channels with fast (α1) and slow (α6) kinetics and GABA spillover generating the transient and sustained components of inhibition observed experimentally. In order to account for experimental results ^11^, the parameters describing presynaptic dynamics were: release probability = 0.35, τREC = 36 ms, τfacil = 58.5 ms and τI = 0.1 ms, respectively ^31^.

The GoC-GoC synapses was modeled as a GABAergic dual exponential synapse and fitted to available data ^3^; rise time and decay time constants were set to 0.9 ms and 10 ms respectively, while maximal conductance to 130 pS.

The gap junctions are electrical synapses among GoCs dendrites. In absence of external input, gap junctions promote GoCs synchronization; sparse MFs input can determine both excitatory and inhibitory effects ^5^. In the present model, a maximal conductance of 50 pS was set for all gap junctions

***Experimental validation against existing data***

Simulations were in close matching with previous experimental data in cerebellar slices. The number of spikes in the center was between 1 and 5, decreasing from core to periphery, in agreement with quantitative estimates using simultaneous patch-clamp and calcium imaging ^32^. Moreover, the diameter of the responding area (~ 100 μm) was quite similar to values measured using multielectrode local field potential recordings ^33^ and voltage-sensitive dye imaging ^34^. Interestingly, the number of GrCs that responded with spikes (here 330) approached the number estimated from local field potential reconvolution in response to tactile stimulation *in vivo* (~500) ^35^. These results altogether suggest that the granular layer can operate through the activation of microcircuit units roughly corresponding to activation of ~50 contiguous GLOMs. In the model, a MF bundle activating 48 adjacent GLOMs contacted 1378 GrCs and 23 GoCs and could influence an extended GoCs network indirectly through gap junctions, PFs and reciprocal inhibitory synapses. Activation of this MF bundle with a spike burst ^21,25, 36, 37^ caused about 300 of these GrCs to generate action potentials. This figure corresponds to the dense response clusters elicited *in vivo* by punctuate tactile facial stimulation, that were estimated to contain about 250 spiking GrCs ^35^.

***MODELING OF LONG-TERM SYNAPTIC PLASTICITY***

Plasticity was modelled following the approach described in ^38^and ^39^. The equations used below have been adapted mainly from ^38^. According to the Ca^2+^ control hypothesis ^40^, synaptic plasticity is a non-linear [BCM-shaped] function of the activity-dependent rise in postsynaptic Ca^2+^ concentration: large Ca^2+^ transient should lead to LTP , while a moderate increase in Ca^2+^ concentration determines LTD. When the Ca^2+^ concentration change is negligible or null, no synaptic plasticity of any kind takes place. The computational model proposed by Shouval for the hippocampus has been adapted to the mossy fibers - granule cell synapse as follows.

***Ca^2+^ current though NMDA receptor***

The NMDA Ca^2+^ current, as a function of membrane potential, is modeled by the equation:

$$I_{NMDA(t_{i})}= P_{0}G_{NMDA}\left[ I_{f}\theta\left( t_{i} \right)e^{{-t_{i}}/{\tau_{f}}}+ I_{s}\theta\left( t_{i} \right)e^{{-t_{i}}/{\tau_{s}}} \right]B(V)(V- V_{r})$$

with glutamate binding occurring at t = 0.

*I­_f_­­* and *I_s_* represent slow and fast components of NMDA receptor current, respectively. In our model, *I_f_* = 0.35 and *I_s_* = 0.65. The time constants τ_f_ and τ_s_ have been set to 50 ms and 200 ms.

*θ* is a function equal to zero if its argument is negative and one if its argument is positive.

*G_NMDA_* , the receptor conductance, is estimated to be -1/53 [μM/(ms * mV)] ^41-45^.

*P_0_* is the fraction of NMDARs moving from closed to open state after a presynaptic spike. Previous experimental and modeling studies showed that NMDA channels are characterized by a low channel open probability ^14, 45^; in this work, *P_0_* is equal to 0.1425

The term *B(V)(V - Vr)* describes the voltage (*V*) dependence of the Ca^2+^ current; (*V-Vr*) represents the driving-force, with *Vr* = 130 mV being the Ca^2+^ reversal potential.

Magnesium block is expressed by *B* :

$$B= \frac{1}{1+\exp-(\frac{V-V_{0}}{k})}$$

with k = -13 mV and V_0_ = -20 mV (see Nieus et al., 2006 for details). Different parameter models may be tested in the future to verify their impact on plasticity (e.g. see ^46^)

***Ca^2+^ concentration***

To calculate Ca^2+^ concentration as a function of Ca^2+^ current through NMDARs (*I_NMDA_)* in postsynaptic cell, the following equation has been used:

$$\frac{d[Ca\left( t \right)]}{dt} =I_{NMDA}\left( t \right)-(1/{\tau_{Ca})[Ca\left( t \right)]}$$

where *[Ca(t)]* is the Ca^2+^ concentration at time *t* and *τ_Ca_* is the decay time constant of calcium. In all simulations, *τ_Ca_* = 150 ms, according to previous experimental work ^47^

***The* Ω *function***

The Ω function expresses the relation between Ca^2+^ concentration and changes in synaptic weight (W).

Ω *= 0.5 + sig(Ca – α2, β2) – 0.5 sig(Ca – α1, β1)*

Where *sig* is a sigmoidal function defined as:

*sig(x, β) = exp(βx) / (1 + exp(βx))*

with *α1* = 0.25, *α2* = 0.77, *β1* = 80 and *β2* = 80. These values allowed to fit the plasticity model to existing experimental data ^48^.

***The η function***

The *η* function represents the calcium-dependent learning rate and is inversely related to the learning time constant *τ* such that *η = 1/τ* , with:

$$\tau= \frac{P_{1}}{P_{2}+{(Ca)}^{P_{3}}}+P_{4}$$

The values of parameters have been taken from Rackham et al., 2010: P_1_ = 100 ms, P_2_ = 0.002, P_3_ = 4, P_4_ = 1000 msec.

***Synaptic weight change after plasticity***

The change in synaptic strength was calculated as follows:

*W_post_ =* η*([Ca])(*Ω*([Ca]) – W_pre_)*

Where *W_pre_* = weight before plasticity and *W_post_* = weight after plasticity.

*W_pre_* is equal to 0.5. Therefore, when *W_post_* > 0.5 there will be LTP, when *W_post_* < 0.5 LTD instead.

In this work, plasticity was implemented through release probability (*P*) variation: before plasticity, we have *P_pre_* equal to 0.42 for all MF-GrCs synapses. Therefore, *P_post_* can be estimated from *W* variation as follows:

$$P_{post}= \frac{P_{pre} \times W_{post}}{W_{pre}}$$

For example, if *W_post_* = 0.8 then *P_post_* = (0.42*0.8) / 0.5 = 0.672, which is LTP; if *W_pos_*_t_ = 0.2, *P_post_* = 0.168, that is LTD.

***SUPPLEMENTAL FIGURES***

***Fig. s1. Anatomical connectivity in the granular layer network model.*** (A) Scheme of the granular layer network and circuit wiring reconstructed in the model. The granule cells are activated by MFs terminating in the GLOMs and are inhibited by GoCs. The GoCs are excited through MFs, AAs and PFs, are coupled through gap junctions and are inhibited by reciprocal inhibitory synapses. The network activity was elicited by stimulating the GLOMs. The full network reproduces a 800 x 800 x 150 µm^3^ portion of the granular layer, comprising 484000 GrCs, 915 GoCs and 29415 GLOMs. Graphic rendering is obtained using Python visualization tools. (B) Schematics of granular layer wiring between GrCs, GoCs and GLOMs. (C) Schematics of wiring between GoCs.

***Fig. s2. Anatomical connectivity in a granular layer model response unit.*** (A) Localization and number of neurons in a unit. Active GLOMs (n=42), Excited GrCs (n=300), Excited GoCs (by MFs n= 10, by AAs n= 30). (B) Number of excited and inhibited dendrites per GrC in the unit. Note that the higher density of excitatory and inhibitory connections is in the core of the recruited volume.

***Fig. s3. Spike discharge properties in a granular layer model response unit.*** The panels show the number of spikes and the first spike delay in the granular layer response unit of Fig. S2. The parameters are represented in color in two conditions: inhibition ON (control condition) and inhibition OFF (imitating the experimental application of a GABA receptor inhibitor like gabazine). Note that:

- When inhibition is ON, the network fine tunes the distribution of first spike delays over 0-50 ms but much less so the number of spikes, which is limited by feed-back inhibition (most cell generate just 1-2 spikes). When inhibition is OFF, the opposite occurs: the network fine tunes the number of spikes between 0-20 per cell but much less so the distribution of first spike delays, with most cells discharging after a few ms.
- The response unit closely resembles the «Dense Cluster» activated in the granular layer in vivo and inferred by model based LFPs reconvolution ^35^.

***Fig. s4. Subcellular mechanisms in granule cells during plasticity.*** The traces show the membrane voltage, AMPA current, NMDA current and GAGA-A current before and after plasticity for two granule cells, one located in the core and the other in the periphery. Note potentiation of firing in the core cell but absence of potentiation in the peripheral cell.

***Fig. s5. Regulation of plasticity in the response unit by granular layer synapses.*** (A) Effect f the specific switch off of the mechanisms regulating synaptic inhibition. As explained in the main text, the full switch off of inhibition had a dramatic effect on plasticity. However, the LTP/LTD balance was not much changed by the synaptic mechanisms of the granular layer network ^2, 28^ taken individually. This suggest that these mechanisms may be instrumental to regulate microcircuit dynamics more than mossy fiber - granule cell plasticity. (B) Release probability changes with synaptic inhibition ON and OFF.

***SUPPLEMENTAL REFERENCES***

10.	Eccles, J.C., Ito, M. & Szentagothai, J. *The cerebellum as a neural machine* (Springer-Verlag., Berlin, Heidelberg, New York, 1967).
