## Supplementary figures and images for "Cellular-resolution mapping uncovers spatial adaptive filtering at the cerebellum input stage"

### Supplemental figure 1

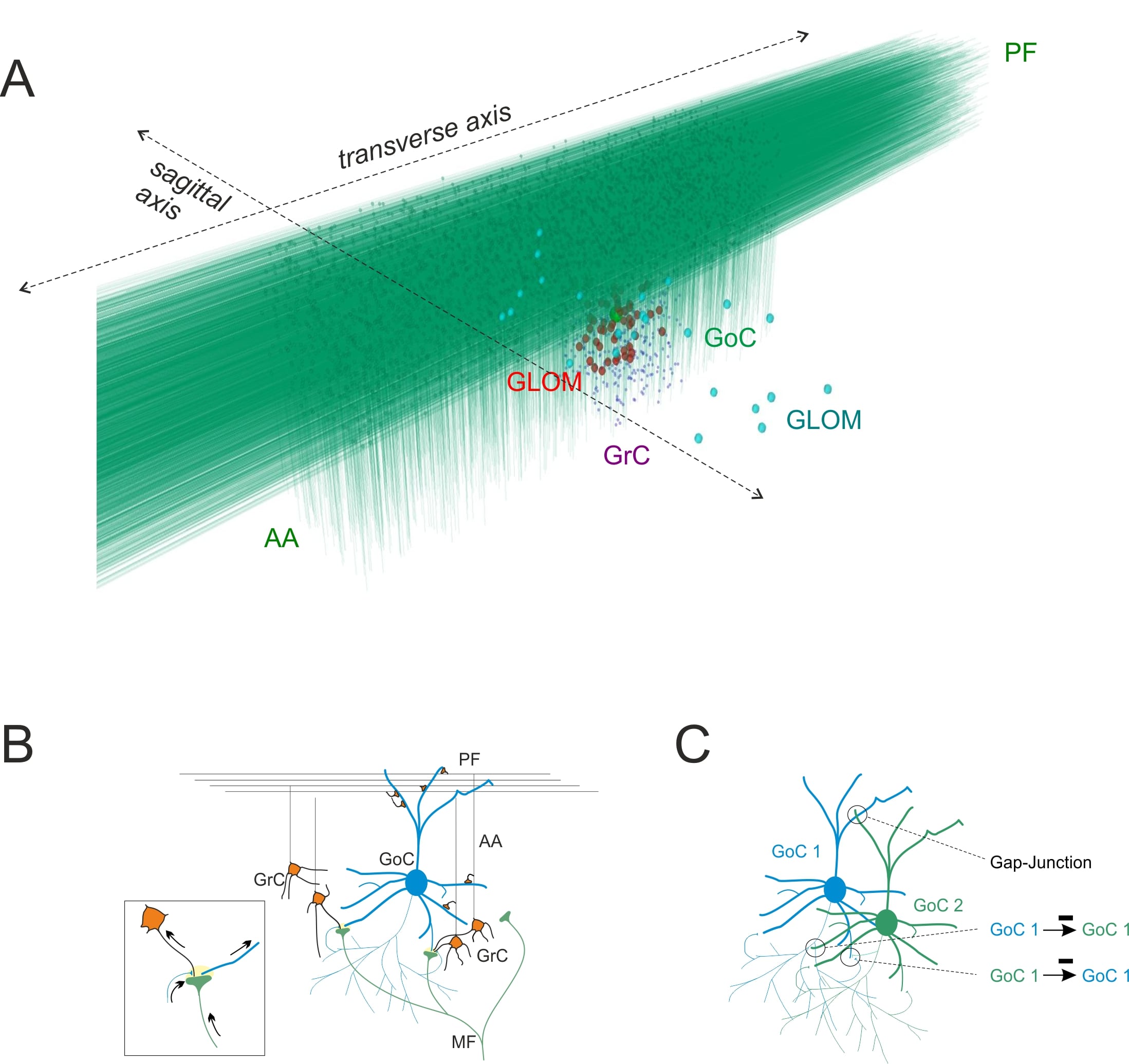

### Supplemental figure 2

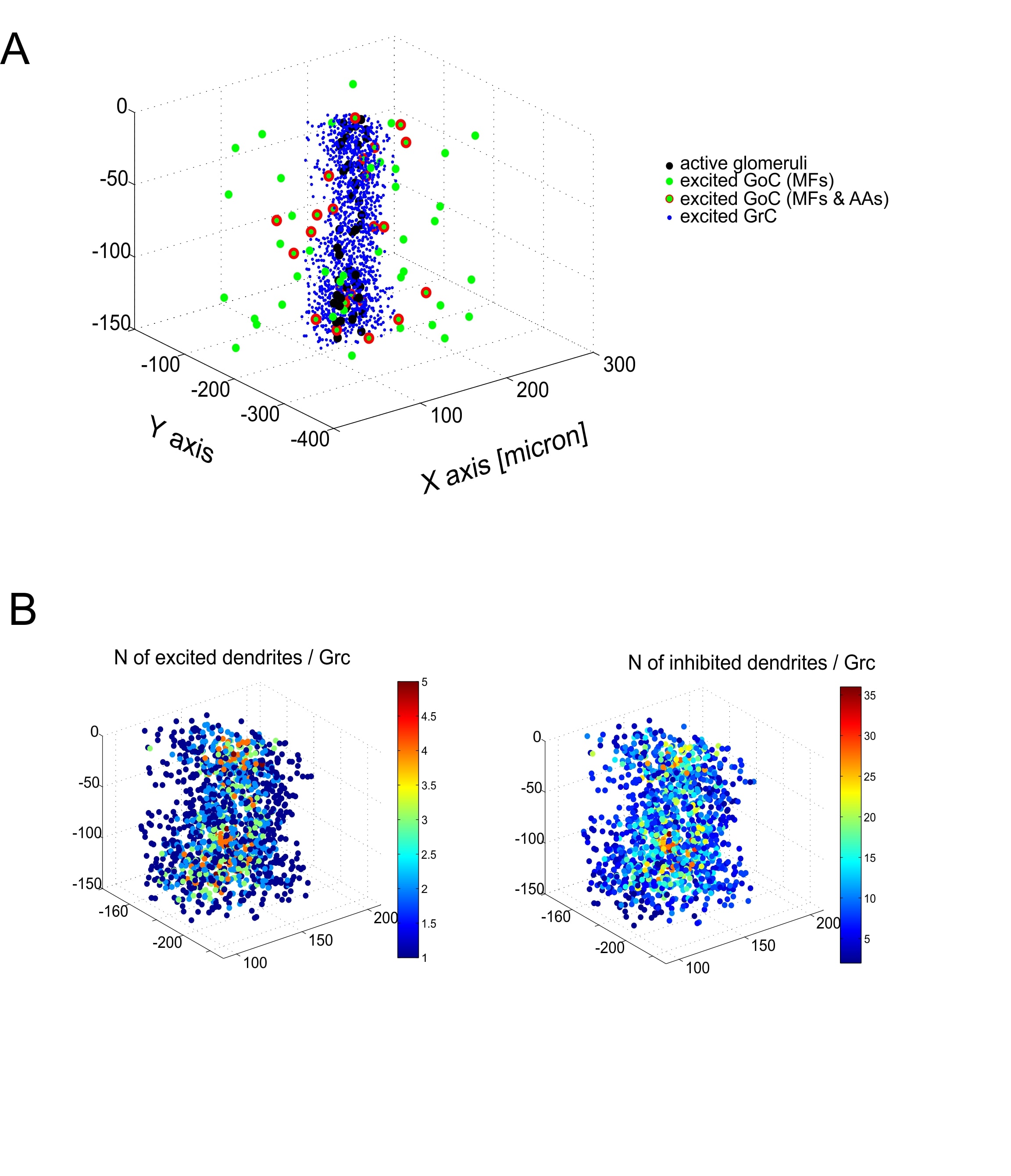

### Supplemental figure 3

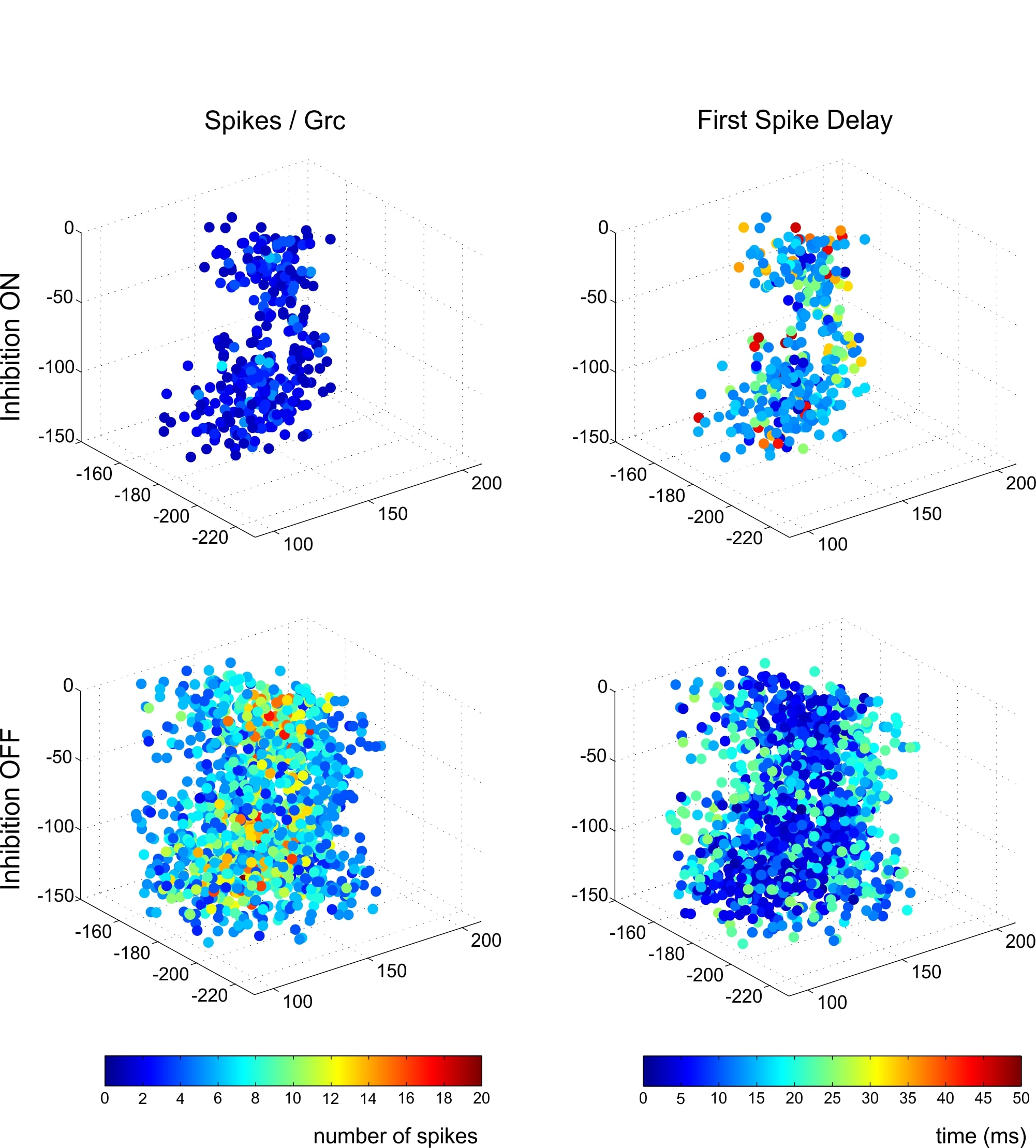

### Supplemental figure 4

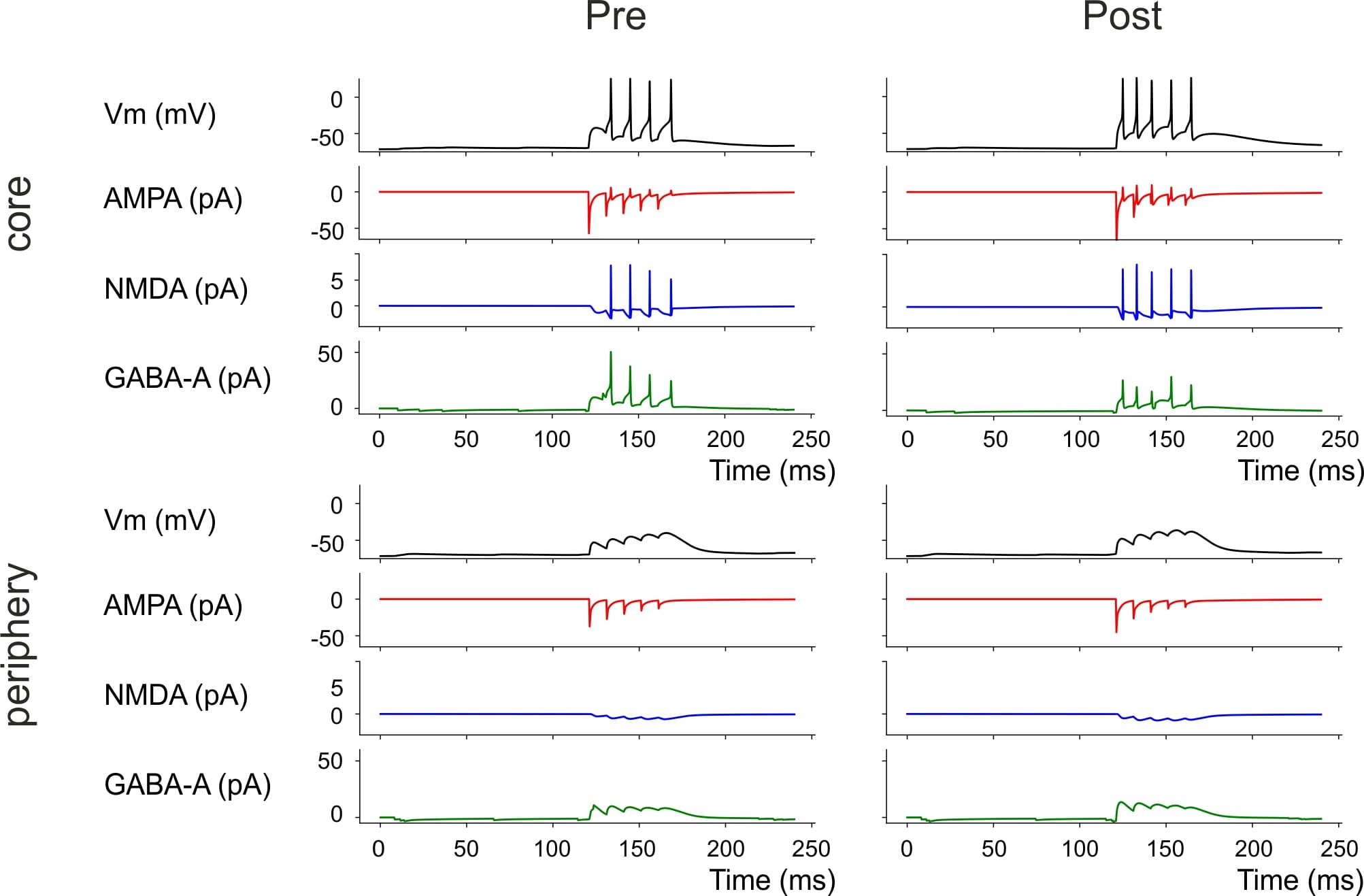

### Supplemental figure 5

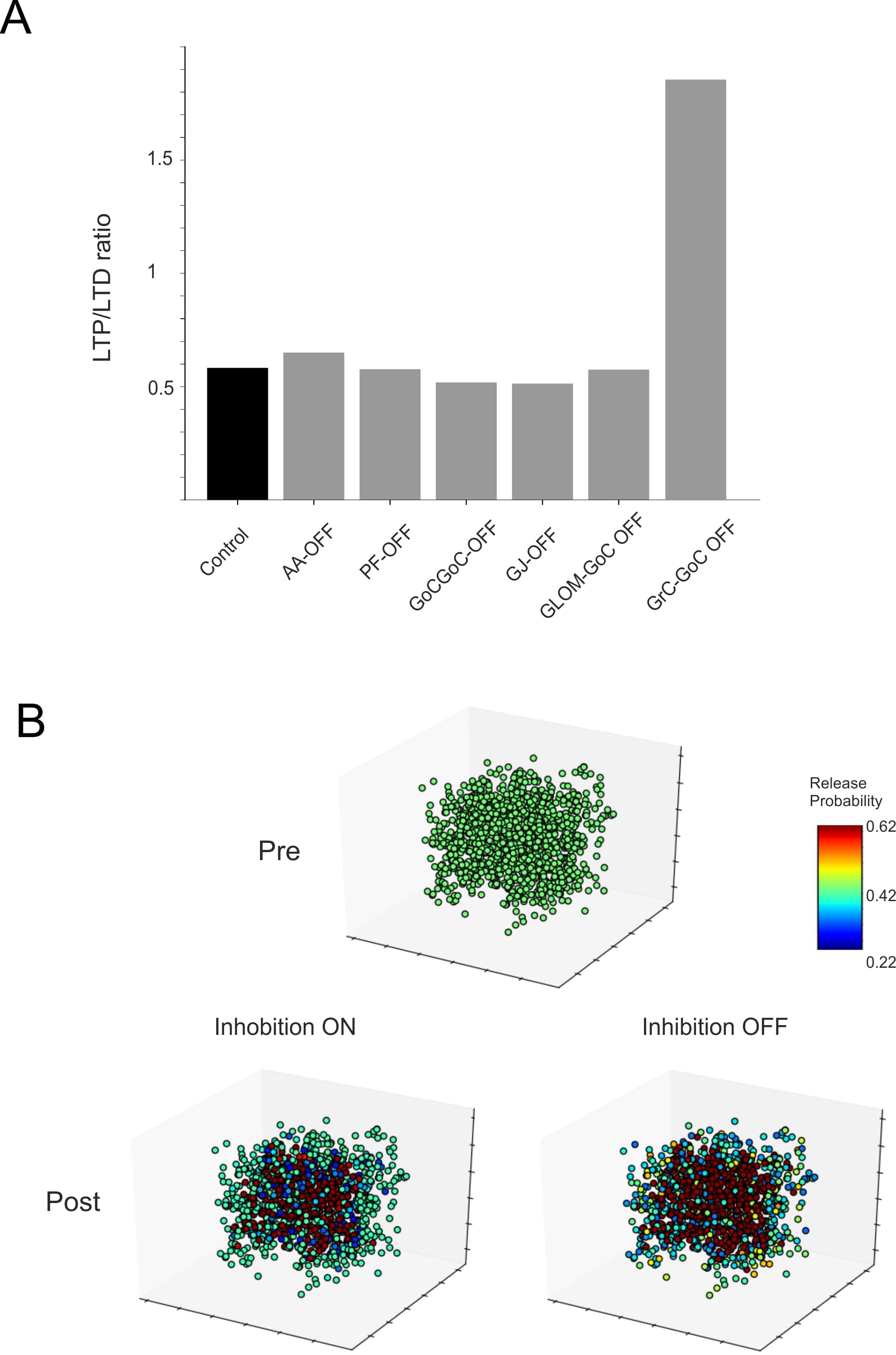
